# Melatonin mitigates oxidative stress and metabolic dysfunction induced by interleukin-6 and dopamine in SH-SY5Y cells

**DOI:** 10.1101/2025.09.03.673920

**Authors:** Heather K. Macpherson, Roger B. Varela, Venea Dara Daygon, James P. Kesby, Xiaoying Cui, Susannah J. Tye

## Abstract

Melatonin has emerged as a promising pharmacological candidate for bipolar disorder (BD), though its mechanisms of action remain incompletely understood. Its antioxidant, anti-inflammatory, and anti-dopaminergic properties suggest potential relevance to BD pathophysiology. This study investigated melatonin’s effects on dopamine signalling, metabolism, and oxidative stress under inflammatory and hyperdopaminergic conditions in differentiated SH-SY5Y neuronal cells. Cells were pretreated with 100nM melatonin or vehicle for 2 hours, then exposed to vehicle, IL-6 (20ng/mL), dopamine (5µM or 500µM), or dopamine (500µM) with ascorbic acid (1mM) for 12 or 24 hours. Dopaminergic markers were assessed via real-time PCR and HPLC; metabolic outcomes were measured using Seahorse assay, central carbon metabolomics, in-cell Western assay, and glucose uptake assay; and oxidative stress was evaluated via reactive oxygen species (ROS), superoxide (SOX), and total antioxidant capacity (TAC) assays. IL-6 increased dopamine levels, p-Erk1/2/Erk1/2, p-AMPK/AMPK, nucleotide pools, and TAC, while reducing dopamine turnover, SV2C expression, and spare respiratory capacity. Melatonin alone increased nucleotides and NADH, while reducing dopamine turnover, ROS, and glucose-1-phosphate. In IL-6 conditions, melatonin pretreatment enhanced spare respiratory capacity, glucose uptake, and NADH, while reducing dopamine, TAC, p-AMPK/AMPK, p-GSK3β/GSK3β, and non-mitochondrial oxygen consumption. High-dose dopamine (500µM) elevated SOX, p-Erk1/2/Erk1/2, insulin receptor-α, GLUT1, glycolytic ATP (glycoATP), and non-mitochondrial oxygen consumption. Melatonin pretreatment attenuated p-Erk1/2/Erk1/2 and GLUT1 elevations. Combined dopamine and ascorbic acid further increased glycolytic intermediates, ROS, p-AMPK/AMPK, and TAC, while reducing p-Erk1/2/Erk1/2, p-mTOR, GLUT1, glucose uptake, and glycoATP. Overall, melatonin mitigated IL-6-induced dopaminergic, oxidative, and metabolic alterations, and partially protected against dopamine-induced metabolic shifts. These findings suggest melatonin may alleviate manic symptoms in BD via both direct dopaminergic modulation and indirect antioxidant and metabolic regulatory effects.

## 1 Introduction

Bipolar disorder (BD) is a complex psychiatric illness characterised by episodic mood fluctuations between mania and depression. In addition to these psychiatric symptoms, BD is associated with widespread physiological changes, including dopamine dysregulation, systemic inflammation, oxidative stress, and impaired cellular metabolism (Berk et al., 2007; Madireddy & Madireddy, 2022). These disturbances are thought to contribute to comorbid conditions such as obesity, type 2 diabetes, and metabolic syndrome (Berk et al., 2007; Madireddy & Madireddy, 2022; Miola et al., 2022). Current treatment strategies for BD are predominantly pharmacological. However, these are often only partially effective and may exacerbate physical health issues (Nierenberg et al., 2023). Consequently, there is growing interest in identifying novel therapies or repurposing existing agents that can address both psychiatric and physiological domains of BD (Geddes & Miklowitz, 2013).

Melatonin has emerged as a promising therapeutic candidate for BD. Both primary and adjunctive melatonin therapies have demonstrated clinical efficacy in reducing symptoms during manic and depressive episodes (Bersani & Garavini, 2000; Livianos, Sierra, Arques, Garcia, & Rojo, 2012; Moghaddam et al., 2020; Nierenberg, 2009; Quested et al., 2021). However, its mechanisms of action remain incompletely understood. A growing body of evidence suggests that hypomelatoninaemia may play a significant role in BD pathophysiology. Patients with BD exhibit reduced plasma melatonin levels across mood states (Bradley et al., 2017; Kennedy, Kutcher, Ralevski, & Brown, 1996; Lam et al., 1990; Robillard et al., 2013), which may reflect impaired melatonin synthesis, increased catabolism, or heightened sensitivity to light-induced melatonin suppression (Dmitrzak- Weglarz et al., 2021; Eslami Amirabadi et al., 2015; Etain et al., 2012; Fan et al., 2010; Lewy et al., 1985; Lewy, Wehr, Goodwin, Newsome, & Rosenthal, 1981; Liu, Huang, Luo, Wu, & Li, 2016; Nathan, Burrows, & Norman, 1999; Nurnberger et al., 2000; Yang et al., 2021). Melatonin has established antioxidant, anti-inflammatory, and anti-dopaminergic properties (Chitimus et al., 2020; Pe et al., 2025), and may also regulate cellular metabolism (Cardinali & Vigo, 2017). These actions align with core biological disruptions observed in BD (Jiki, Lecour, & Nduhirabandi, 2018; Pe et al., 2025).

Importantly, dopamine dysregulation, inflammation, oxidative stress, and metabolic dysfunction are interconnected. High dopamine levels can promote oxidative stress via auto-oxidation and reactive oxygen species (ROS) production (Meiser, Weindl, & Hiller, 2013).

Inflammation exacerbates oxidative stress and alters dopamine signalling (Felger & Treadway, 2017; Hussain et al., 2016), while mitochondrial dysfunction disrupts energy production and increases ROS (Kageyama et al., 2025). Fluctuations in dopamine and energy homeostasis may underlie mood disturbance in BD, with hyperdopaminergia and high energy underpinning manic episodes, and hypodopaminergia and low energy contributing to depressive episodes (Ashok et al., 2017; Bakshi & Kelley, 1991; Baldo, Sadeghian, Basso, & Kelley, 2002; Berk et al., 2007; Morris et al., 2017; Scott & McClung, 2023; Winter et al., 2007).

In line with this, patients with BD demonstrate increased plasma, serum, brain, and cerebrospinal fluid markers of oxidative stress and decreased levels of antioxidants when compared to healthy controls (Y. Kim et al., 2019). Chronic inflammation is also considered to be a trait marker of BD, with patients in all mood states showing significant increases in inflammatory markers (Madireddy & Madireddy, 2022). Interleukin-6 (IL-6) is of particular interest as a putative biomarker of BD, as it is consistently elevated in euthymic, manic, depressed, and mixed mood states (Barbosa et al., 2013; Brunoni et al., 2020; Y. K. Kim, Jung, Myint, Kim, & Park, 2007; Koga et al., 2019; Pandey, Ren, Rizavi, & Zhang, 2015; Uyanik, Tuglu, Gorgulu, Kunduracilar, & Uyanik, 2015; Vasconcelos-Moreno et al., 2017).

Investigating the role of melatonin in modulating oxidative stress, metabolism, and dopamine in high inflammation and high dopamine environments may provide much-needed insight into the mechanisms underpinning melatonin treatment in BD, as well as the overall aetiology of the disease. This study aimed to determine whether melatonin could attenuate the molecular sequelae of inflammation and elevated dopamine in differentiated SH-SY5Y cells, which is a commonly-used and validated model of mature, human catecholaminergic neurons (Kovalevich & Langford, 2013). Specifically, we assessed the effects of melatonin on oxidative stress, metabolism, and dopamine-related markers following exposure to IL-6 or dopamine. We also compared melatonin’s effects to those of ascorbic acid, to determine whether antioxidant activity is a key mechanism of action.

## 2 Methods

### 2.1 Experimental design

Two separate studies are described in this article. The first study aimed to examine the effects of IL-6 and/or melatonin on dopaminergic, metabolic, and oxidative stress markers using quantitative real-time PCR (RT-PCR), high-performance liquid chromatography (HPLC), in-cell Western assays (ICWs), carbon central metabolite (CCM) metabolomics, glucose uptake assay, Seahorse assay, reactive oxygen species (ROS) / superoxide (SOX) assay, and total antioxidant capacity (TAC) assay. The second study aimed to examine the effects of dopamine either alone or in combination with melatonin or ascorbic acid on metabolic markers using ICWs, CCM metabolomics, glucose uptake assay, Seahorse assay, ROS/SOX assay, and TAC assay. Details (including brands and catalogue numbers) of all reagents, kits, software, equipment, and instruments used in this study can be found in Supplementary Data 1.

### 2.2 Cell culture and differentiation

SH-SY5Y cells were chosen for this experiment as they exhibit a mature neuron-like phenotype when differentiated for at least three days with 10µM retinoic acid (RA) (Kovalevich & Langford, 2013), and are frequently used for investigations of dopamine function. SH-SY5Y cells are not listed as a commonly misidentified cell line by the International Cell Line Authentication Committee, according to the most recent version of the database (Capes-Davis et al., 2010). This cell line was validated as possessing dopaminergic properties post-differentiation (including a significant increase in tyrosine hydroxylase (TH) when compared to undifferentiated cells). SH-SY5Y cells were cultured in Dulbecco’s Modified Eagle Medium/Nutrient Mixture F-12 (DMEM/F12), 10% heat-inactivated foetal bovine serum (FBS), 1% GlutaMAX™ 100X, and 100 units/ml penicillin-streptomycin, and incubated at 37°C, 5% CO_2_. Cells were passaged 23-26 times before use. Cells were seeded on 24-well plates coated with 0.1mg/ml poly-D-lysine at a density of 8 x 10^4^ cells per well for RT-PCR, HPLC, TAC, and CCM metabolomics; on uncoated 96-well plates at a density of 6 x 10^3^ cells per well for ICW, glucose uptake, and ROS/SOX assays; and on uncoated 96-well Seahorse XFe96 Pro Cell Culture Microplates at a density of 3 x 10^3^ cells per well for the Seahorse assay. Cells were then treated with differentiation medium (DMEM/F12, 1% heat-inactivated FBS, 1% GlutaMAX™ 100X, 100 units/ml penicillin-streptomycin, 2% B27™ Supplement 50X, and 10µM all-trans RA) for seven days. This medium was changed every two to three days.

### 2.3 Treatments

The drugs used in this experiment included the following: 100nM melatonin dissolved in 0.001% ethanol and 0.001% DMSO, 20ng/ml IL-6, 5µM and 500µM dopamine, and 1mM L-ascorbic acid. Cells that weren’t treated with melatonin were supplemented with vehicle (0.001% ethanol, 0.001% DMSO). After seven days of differentiation, cell media was replaced with RA-free differentiation media. Melatonin was applied two hours prior to other treatments; cells were then treated for twelve hours (for Experiment 1 RT-PCR and all Experiment 2 assays) or twenty-four hours (for all other Experiment 1 assays).

### 2.4 Quantification of dopaminergic markers

#### 2.4.1 Gene expression analysis

RT-PCR was used to quantify the expression of melatonin receptor genes, and genes related to dopamine synthesis, transport, release, and metabolism. RNA was extracted from treated cells (six biological replicates per group) using the RNeasy^®^ Mini kit and the RNAse-free DNAse kit according to the manufacturer’s instructions. RNA concentrations were measured using the NanoPhotometer N60 microvolume spectrophotometer. 800ng total RNA per sample was used for cDNA synthesis. cDNA was synthesised from total RNA using the SensiFAST™ cDNA synthesis kit according to the manufacturer’s instructions. RT-PCR was performed using the SensiFAST™ SYBR^®^ No-ROX kit according to the manufacturer’s instructions. The following primers were used: HPRT (used as endogenous control), TH (tyrosine hydroxylase), MAOA (monoamine oxidase A), COMT (catechol-O-methyltransferase), VMAT2 (vesicular monoamine transporter 2), SV2C (synaptic vesicle glycoprotein 2C), MTNR1A (melatonin receptor 1), and MTNR1B (melatonin receptor 2) (see Supplementary Data 2 for primer sequences). RT-PCR consisted of a denaturation step (90°C for 10 minutes), then amplification (95°C for 10s, 60°C for 30s and 72°C for 20s) for 45 cycles, performed in the LightCycler^®^ 480 Instrument II. The ΔCt method was used to assess gene expression.

#### 2.4.2 High-performance liquid chromatography (HPLC)

HPLC with electrochemical detection was used to measure dopaminergic markers (dopamine and homovanillic acid (HVA)), serotonin, 5-hydroxyindoleacetic acid, and noradrenaline). Cells (six biological replicates per group) were homogenised in 0.1M perchloric acid containing 50 ng/ml deoxyepinephrine (internal standard) using a probe sonicator, centrifuged at 4°C for five minutes at 10000rpm, and filtered using a 22µM nylon syringe filter. 30µl of each sample was injected at a flow rate of 0.6ml/min into the HPLC system, which consisted of an autosampler, an isocratic HPLC pump, a Coulochem III electrochemical detector, and a SunFire C8 3µm, 4.6 x 100 mm column. The mobile phase was composed of 12% acetonitrile, 50mM citric acid, and 25mM potassium dihydrogen phosphate, with 1.4 mM octane sulfonic acid and 1mM ethylenediaminetetraacetic acid (EDTA), adjusted to pH 4.3 with phosphoric acid. Detection was performed using a Model 5014B Microdialysis High Performance Analytical Cell, with the first and second electrodes maintained at −150 mV and +300 mV, respectively. Data were stored and processed using Chemstation 7.2, and metabolite quantification was achieved by calculating peak-area ratios relative to the internal standard, expressed as ng/ml.

### 2.5 Metabolic assays

#### 2.5.1 In-cell Western assays

ICWs were performed to measure markers of metabolic stress, glycolysis, glucose uptake, and insulin sensitivity. Cells (six biological replicates per group) were incubated in ice-cold methanol for twenty minutes, washed thrice with Tris-buffered saline (TBS) + 0.1% Tween-20, then blocked in Intercept Blocking Buffer for 1.5 hours. Cells were then incubated in primary antibodies (see Supplementary Data 3 for details) diluted in Intercept Blocking Buffer + 0.2% Tween-20 overnight at 4°C under gentle agitation. Cells were washed four times and then incubated in IRDye 800CW Goat anti-rabbit IgG secondary antibody (1:500) with Cell Tag 700 Stain (1:500) for one hour at room temperature under gentle agitation in the dark. Cells were washed four times and then imaged on the Odyssey CLx Imaging system at 169µM. Values were standardised according to Cell Tag 700 results.

#### 2.5.2 Central carbon metabolite (CCM) analysis

CCM analysis was used to quantify the levels of metabolites involved in cellular respiration (see Supplementary Data 4 for full list of compounds). Cells (six biological replicates per group) were collected in 1ml 50% ice cold methanol and stored at −80°C until extraction. On the day of extraction, the samples were sonicated in ice water for five minutes; 300µl chloroform was then added. The samples were vortexed, sonicated in ice water for five minutes, and then centrifuged at 16000 x g for five minutes at 4°C. The top aqueous layer was then removed and the methanol evaporated using a vacuum concentrator until only the water fraction was left. This was then freeze dried overnight and reconstituted in 100µl 2% acetonitrile solution. Finally, the samples were centrifuged at 16000 x g for five minutes at 4°C. All standards were serially diluted from 200µM to 1.5nM and added with 5µM azidothymidine. CCM levels were quantified using a previously described liquid chromatography-tandem mass spectrometry (LC-MS/MS) method (Espinosa et al., 2020).

LC-MS/MS was performed using a Nexera ultra high-performance liquid chromatograph coupled to the 8060 triple quadrupole mass spectrometer on Acquity HSS T3 1.8μm, 2.1mm x 100mm columns with Acquity HSS T3 1.8µm 2.1mm x 5mm guard columns. LC-MS/MS was performed on negative ionisation mode with a 250µl/min flow rate, 10µl injection volume, and column temperature of 45°C. 7.5mM tritbutylamine with acetic acid added until the solution reached pH 4.95 was used for mobile phase A, and 100% acetonitrile for mobile phase B. Total analysis time per sample was 35 minutes.

#### 2.5.3 Glucose uptake assay

Glucose uptake was measured using a commercially available kit according to the manufacturer’s instructions. Cells (six biological replicates per group) were first washed with phosphate-buffered saline (PBS); 1mM 2-deoxyglucose was added, and the cells were incubated for fifteen minutes at room temperature. Stop solution and neutralisation solution were added, with a thirty second gentle agitation step after each addition. Cells were then incubated in 2-deoxyglucose-6-phosphate detection reagent for one hour at room temperature. Luminescence was measured using the CLARIOstar Plus microplate reader.

#### 2.5.4 Seahorse assay

Mitochondrial respiratory function and bioenergetic health were measured using the Seahorse XF Cell Mito Stress Test assay according to the manufacturer’s instructions. Cells (nine biological replicates per group) were treated as usual. At the end of the treatment period, differentiation media was aspirated and 180µl Seahorse XF DMEM supplemented with 2mM glutamine, 1mM pyruvate, and 10mM D-glucose was added to each well. Cells were then incubated at 37°C in a CO_2_-free environment for 45 minutes. The Mito Stress assay was performed in the Seahorse XFp Analyzer using 2.5µM oligomycin in port A, 1.0µM carbonyl cyanide-p-trifluoromethoxy phenylhydrazone (FCCP) in port B, and 0.5µM rotenone/antimycin in port C. Readings were obtained in triplicate. Agilent Seahorse Analytics software was used to calculate key assay parameters, including basal respiration, basal glycolytic ATP production (glycoATP), basal mitochondrial ATP production (mitoATP), ATP-linked respiration, proton leak, maximal respiration, non-mitochondrial oxygen consumption, spare respiratory capacity, and coupling efficiency. Values were standardised according to Cell Tag 700 results.

### 2.6 Oxidative stress assays

#### 2.6.1 Cellular reactive oxygen species (ROS) / superoxide (SOX) assay

Cellular ROS/SOX levels were quantified using a commercially available detection assay kit. An hour prior to the end of treatment, cells (ten biological replicates per group) were incubated at room temperature for thirty minutes; a ROS inducer (pyocyanin) was added to control wells, and the ROS/SOX detection solution to all wells. Cells were then incubated at 37°C in the dark for another thirty minutes. Fluorescence was measured using the CLARIOstar Plus microplate reader.

#### 2.6.2 Total antioxidant capacity (TAC) assay

Total antioxidant capacity was quantified using a commercially available kit according to the manufacturer’s instructions. Cells (four biological replicates per group) were harvested from 24-well plates using trypsin-EDTA (0.25%). Samples were then quenched with differentiation media, centrifuged at 500 x g for five minutes at 4°C, washed with cold PBS, centrifuged again, then homogenised in ultrapure water. Cell homogenate samples were incubated on ice for ten minutes and centrifuged at top speed for five minutes at 4°C; supernatant samples were then collected. Samples and standards were added in duplicate to a 96-well plate. Cu^2+^ working solution was added to all wells; the plate was then incubated for ninety minutes at room temperature under gentle agitation in the dark. Absorbance was measured using the CLARIOstar Plus microplate reader.

### 2.7 Statistical analyses

All statistical analyses were performed on GraphPad Prism 10.0.2. Normality of data was assessed using the Shapiro-Wilk and D’Agostino-Pearson tests, and heteroscedasticity with the Brown-Forsythe test. No tests for outliers were performed. Normal data were analysed using one-way analysis of variance (ANOVA) or Welch’s ANOVA (if non-heteroscedastic); non-normal data were analysed using the Kruskal-Wallis test. Post-hoc tests (Holm-Šídák for one-way ANOVA, Dunnett’s T3 for Welch’s ANOVA, and Dunn’s for Kruskal-Wallis) for multiple comparison were performed between vehicle and IL-6, vehicle and melatonin, IL-6 and IL-6 + melatonin, vehicle and 5µM dopamine, vehicle and 500µM dopamine, 500µM dopamine and dopamine + melatonin, 500µM dopamine and dopamine + ascorbic acid, and dopamine + melatonin and dopamine + ascorbic acid groups. The significance threshold was set at p < 0.05 for all statistical tests. Detailed statistical results for post-hoc tests can be found in Supplementary Data 5 and 6.

## 3 Results

### 3.1 Melatonin normalised IL-6-induced hyperdopaminergia in SH-SY5Y cells

Given prior evidence that inflammation disrupts dopamine signalling, we first examined whether IL-6 alters dopamine content or turnover in differentiated SH-SY5Y cells, and whether melatonin could counteract these effects. One-way ANOVA revealed a significant effect of treatment on dopamine content in SH-SY5Y cells [F (3, 17) = 3.274, p = 0.0467]. Post-hoc tests indicated that 20ng/ml IL-6 for 24 hours significantly increased dopamine content when compared to vehicle treatment (Figure 1A). 100nM melatonin treatment alone had no effect on dopamine content, but pre-treatment with melatonin two hours prior to administration of IL-6 significantly suppressed the increase in dopamine content induced by IL-6 (Figure 1A). One-way ANOVA also revealed a trend effect on HVA/dopamine ratio, which is an indicator of dopamine turnover, although results were not statistically significant [F (3, 17) = 2.937, p = 0.0631] (Figure 1B). IL-6 and melatonin both decreased dopamine turnover at a trend level when compared to vehicle treatment (Figure 1B). Furthermore, a trend effect of treatment on SV2C [F (3, 20) = 2.761, p = 0.0689] expression was found (Figure 1C). Post-hoc tests revealed a trend increase in SV2C expression in cells exposed to IL-6 when compared to vehicle (Figure 1C). These results suggest that IL-6 induces a hyperdopaminergic state in neuronal cells and that melatonin can partially suppress this effect, potentially via modulation of dopamine synthesis or clearance mechanisms.

**Figure 1:**
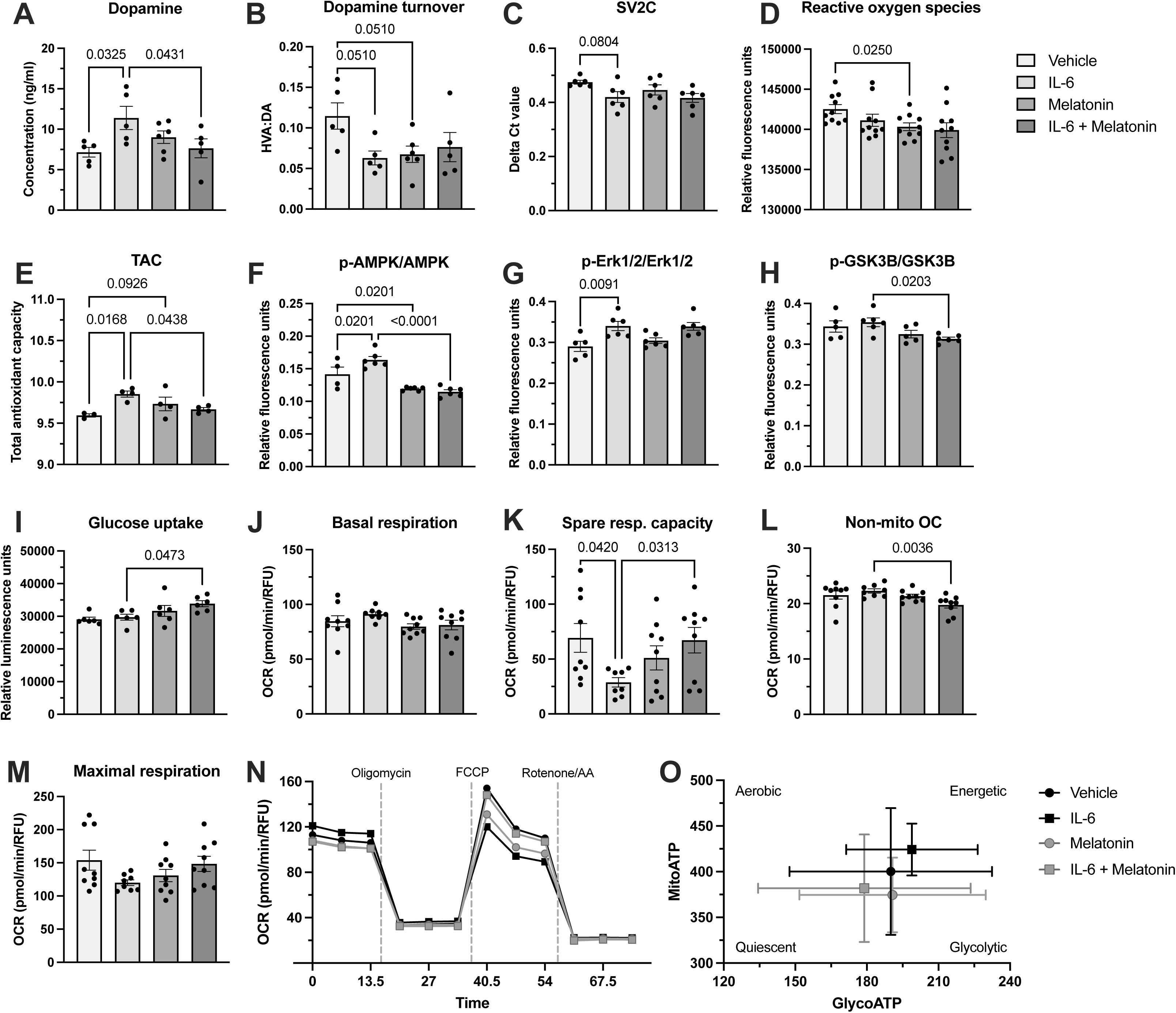
Effects of interleukin-6 (IL-6) and melatonin on dopaminergic, oxidative stress, and metabolic markers in SH-SY5Y cells. Mean (±SEM) concentration (ng/ml) of dopamine (DA) *(A)*, DA turnover (ratio of homovanillic acid (HVA) to DA) *(B)*, gene expression (expressed as ΔCt) of SV2C *(C)*, and levels (relative fluorescence units (RFU)) of reactive oxygen species *(D)*; mean (±SEM) total antioxidant capacity (TAC) *(E)*; mean (±SEM) ratios (RFU) of p-AMPK/AMPK *(F)*, p-Erk1/2/Erk1/2 *(G)*, and p-GSK3β/GSK3β *(H)*; mean (±SEM) levels (relative luminescence units) of glucose uptake *(I)*; mean (±SEM) levels (oxygen consumption rate (OCR) (pmol/min/RFU)) of basal respiration *(J)*, spare respiratory (resp.) capacity *(K)*, non-mitochondrial oxygen consumption (OC) *(L)* and maximal respiration *(M)* compared between vehicle, IL-6, melatonin, and IL-6 + melatonin groups. Effect of treatment is p<0.1 for analyses, as measured by one-way analysis of variance (ANOVA), Welch’s ANOVA, or Kruskal-Wallis test. Displayed p-values reflect results of Holm-Šídák, Dunnett’s T3, or Dunn’s for Kruskal-Wallis post-hoc tests. Seahorse analysis kinetic map of OCR vs. time *(N)* and energetic map of basal mitochondrial ATP production (MitoATP) vs. basal glycolytic ATP production (GlycoATP) *(O)*.

### 3.2 Melatonin reduced oxidative stress in SH-SY5Y cells

As both inflammation and excess dopamine are known to elevate oxidative stress, we next measured ROS, SOX, and TAC levels to determine whether melatonin provides antioxidant protection under these conditions. Welch’s ANOVA revealed a significant effect of treatment on ROS [W (3.000, 19.53) = 3.298, p = 0.0421] (Figure 1D). Melatonin-treated cells demonstrated a significant reduction in ROS levels when compared to vehicle (Figure 1D). A significant effect of treatment was also found on TAC [F (3, 11) = 4.443, p = 0.0281]. When compared to vehicle, IL-6 treatment significantly increased TAC, and a trend towards an increase in TAC was seen with melatonin treatment (Figure 1E). Furthermore, cells treated with IL-6 and melatonin demonstrated significantly lower TAC than cells treated with IL-6 only (Figure 1E). Melatonin significantly reduced ROS and modulated TAC responses, indicating its capacity to restore redox homeostasis—an effect that may contribute to its broader neuroprotective profile.

### 3.3 Melatonin prevented IL-6-induced metabolic stress and mitochondrial dysfunction in SH-SY5Y cells

We then explored whether melatonin modulates IL-6-induced changes in metabolic signalling and mitochondrial function, focusing on markers of metabolic stress, glucose uptake, and bioenergetic health. Significant effects of treatment on p-AMPK/AMPK [F (3, 18) = 20.88, p < 0.0001], p-Erk1/2/Erk1/2 [F (3, 19) = 5.984, p = 0.0048], and p-GSK3β/GSK3β [F (3, 18) = 3.661, p = 0.0321] ratios (Figure 1F-H) were found. IL-6 significantly increased p-AMPK/AMPK ratio, and melatonin significantly decreased p-AMPK/AMPK ratio when compared to vehicle-treated cells (Figure 1F). Furthermore, melatonin pre-treatment prior to administration of IL-6 significantly reduced p-AMPK/AMPK ratio when compared to cells treated with IL-6 alone (Figure 1F). IL-6-treated cells demonstrated a significant increase in p-Erk1/2/Erk1/2 ratio when compared to vehicle-treated cells (Figure 1G). Finally, melatonin pre-treatment prior to IL-6 administration significantly reduced p-GSK3β/GSK3β ratio when compared to IL-6 treated cells (Figure 1H). One-way ANOVA analysis also revealed a significant effect of treatment on glucose uptake [F (3, 20) = 3.748, p = 0.0276] (Figure 1I). Melatonin pre-treatment prior to administration of IL-6 significantly increased glucose uptake when compared to IL-6 treated cells (Figure 1I).

Welch’s ANOVA and Kruskal-Wallis tests revealed significant effects of treatment on basal respiration [W (3.000, 16.60) = 4.524, p = 0.0170], spare respiratory capacity [W (3.000, 15.65) = 5.556, p = 0.0086], and non-mitochondrial oxygen consumption [H (3, 31) = 11.89, p = 0.0078], and a trend effect on maximal respiration [W (3.000, 15.76) = 2.937, p = 0.0656] (Figure 1J-N). IL-6 significantly reduced spare respiratory capacity when compared to vehicle-treated cells (Figure 1K). Melatonin pre-treatment prior to IL-6 administration significantly increased spare respiratory capacity, and significantly reduced non-mitochondrial oxygen consumption when compared to IL-6-treated cells (Figure 1K-L). IL-6 treated cells exhibited a more energetic phenotype than vehicle-treated cells, and cells exposed to melatonin prior to IL-6 exhibited a more quiescent phenotype than IL-6-treated cells (Figure 1O).

Furthermore, a significant effect of treatment on dihydroxyacetone phosphate (DHAP) [F (3, 20) = 5.229, p = 0.0079], glucose 1-phosphate (G1P) [H (3, 20) = 8.367, p = 0.0390], glyceraldehyde 3-phosphate (GA3P) [F (3, 20) = 3.523, p = 0.0338], aconitate [W (3.000, 10.80) = 4.007, p = 0.0381], citrate [F (3, 20) = 3.418, p = 0.0372], fumarate [F (3, 20) = 3.750, p = 0.0275], adenosine monophosphate (AMP) [F (3, 19) = 8.466, p = 0.0009], adenosine diphosphate (ADP) [F (3, 19) = 10.10, p = 0.0003], cytidine monophosphate (CMP) [F (3, 19) = 5.085, p = 0.0094], guanosine monophosphate (GMP) [F (3, 19) = 8.215, p = 0.0010], guanosine diphosphate (GDP) [F (3, 19) = 10.80, p = 0.0002], uridine monophosphate (UMP) [F (3, 19) = 7.268, p = 0.0019], uridine diphosphate (UDP) [F (3, 19) = 9.891, p = 0.0004], and reduced nicotinamide adenine dinucleotide (NADH) [F (3, 18) = 10.82, p = 0.0003], and a trend effect of treatment on acetyl CoA (AcoA) [H (3, 20) = 6.293, p = 0.0982] were found (Figure 2A-O). Significant increases in AMP, ADP, GMP, GDP, UMP, and UDP; a trend increase in CMP; and a significant decrease in ACoA were found in IL-6 treated cells when compared to vehicle-treated cells (Figure 2D, H-N). Melatonin-treated cells also demonstrated increases in AMP, ADP, GMP, GDP, UMP, UDP, and NADH, and a decrease in G1P when compared to vehicle (Figure 2B, H-I, K-O). A significant increase in DHAP and NADH, and trend increases in GA3P, AMP, and GMP were seen in cells pre-treated with melatonin prior to IL-6 administration when compared to cells treated with IL-6 alone (Figure 2A, C, H, K, O). Together, these findings suggest that melatonin not only mitigates IL-6-induced mitochondrial dysfunction, but also improves metabolic efficiency under pro-inflammatory stress.

**Figure 2:**
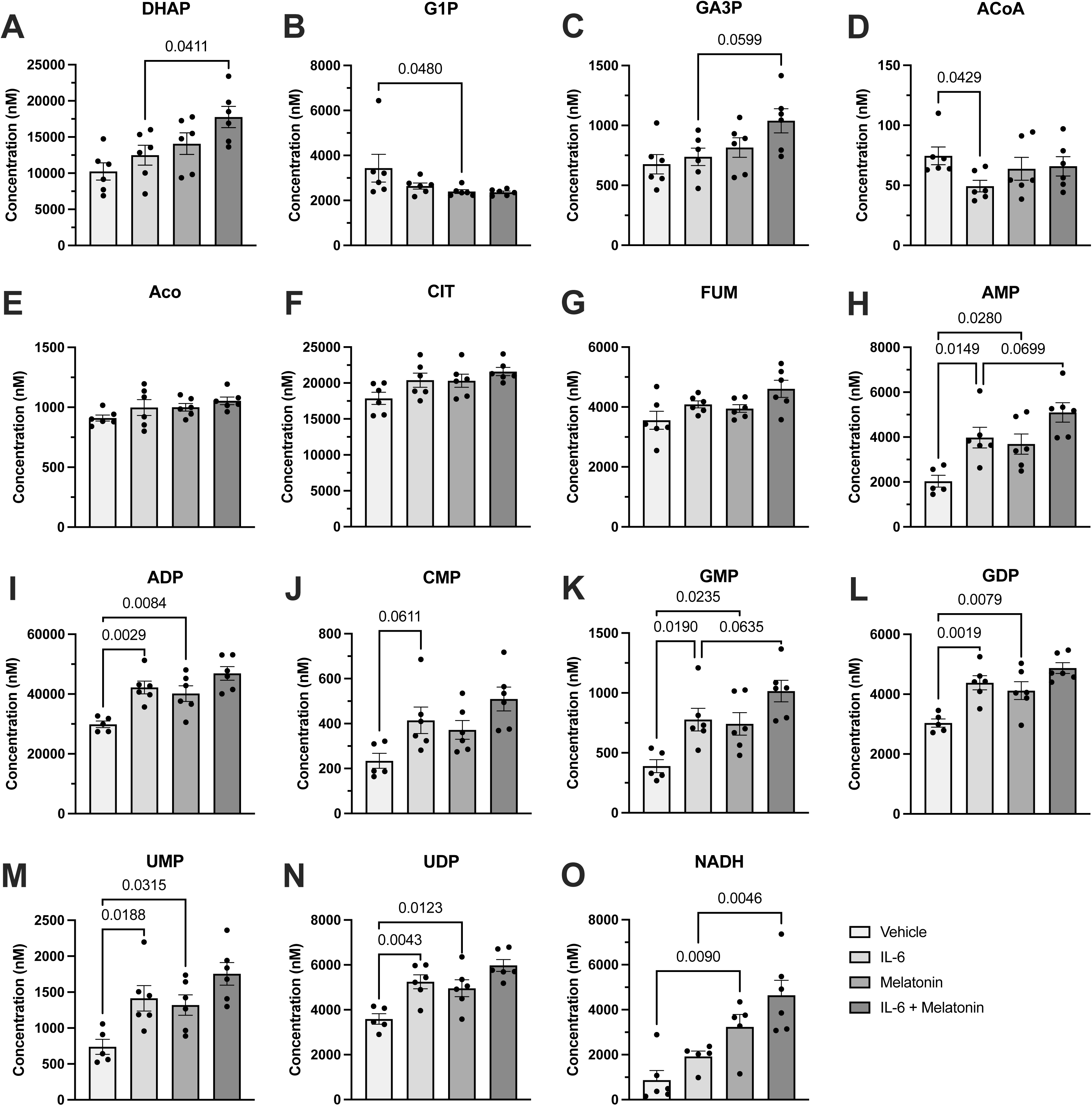
Effects of interleukin-6 (IL-6) and melatonin on central carbon metabolites (CCM) in SH-SY5Y cells. Mean (±SEM) concentration (nM) of dihydroxyacetone phosphate (DHAP) ***(A)***, glucose 1-phosphate (G1P) ***(B)***, glyceraldehyde 3-phosphate (GA3P) ***(C)***, acetyl CoA (ACoA) ***(D)***, aconitate (Aco) ***(E)***, citrate (CIT) ***(F)***, fumarate (FUM) ***(G)***, adenosine monophosphate (AMP) ***(H)***, adenosine diphosphate (ADP) ***(I)***, cytidine monophosphate (CMP) ***(J)***, guanosine monophosphate (GMP) ***(K)***, guanosine diphosphate (GDP) ***(L)***, uridine monophosphate (UMP) ***(M)***, uridine diphosphate (UDP) ***(N)***, and reduced nicotinamide adenine dinucleotide (NADH) ***(O)*** compared between vehicle, IL-6, melatonin, and IL-6 + melatonin groups. Effect of treatment is p<0.1 for analyses, as measured by one-way analysis of variance (ANOVA) or Kruskal-Wallis test. Displayed p-values reflect results of Holm-Šídák or Dunn’s for Kruskal-Wallis post-hoc tests.

### 3.4 500µM dopamine induced changes in oxidative stress markers in SH-SY5Y cells

To determine whether high levels of dopamine independently affect oxidative stress and whether melatonin or ascorbic acid can mitigate its effects, we examined ROS, SOX, and TAC following dopamine administration. One-way ANOVA and Kruskal-Wallis tests revealed a significant effect of treatment on ROS [H (4, 45) = 42.25, p < 0.0001] and SOX [F (4, 45) = 36.03, p < 0.0001] (Figure 3A-B). Cells treated with 500µM dopamine for twelve hours demonstrated a decrease in ROS and an increase in SOX levels (Figure 3A-B). Cells treated with both 500µM dopamine and 1mM ascorbic acid demonstrated an increase in ROS and a decrease in SOX when compared to cells treated with 500µM dopamine alone (Figure 3A-B). Finally, a significant decrease in SOX was seen in cells treated with ascorbic acid and 500µM dopamine when compared to cells treated with melatonin and 500µM dopamine (Figure 3B). A significant effect of treatment was also found on TAC [F (4, 14) = 9.874, p = 0.0005] (Figure 3C). Cells treated with ascorbic acid and 500µM dopamine demonstrated significantly greater TAC than cells treated with 500µM dopamine alone and cells treated with both 500µM dopamine and melatonin (Figure 3C). A trend increase in TAC was also found in cells treated with 500µM dopamine when compared to vehicle (Figure 3C). These results show that interestingly, only ascorbic acid, but not melatonin, reversed dopamine-induced changes in SOX and TAC, suggesting different mechanisms of antioxidant action.

**Figure 3:**
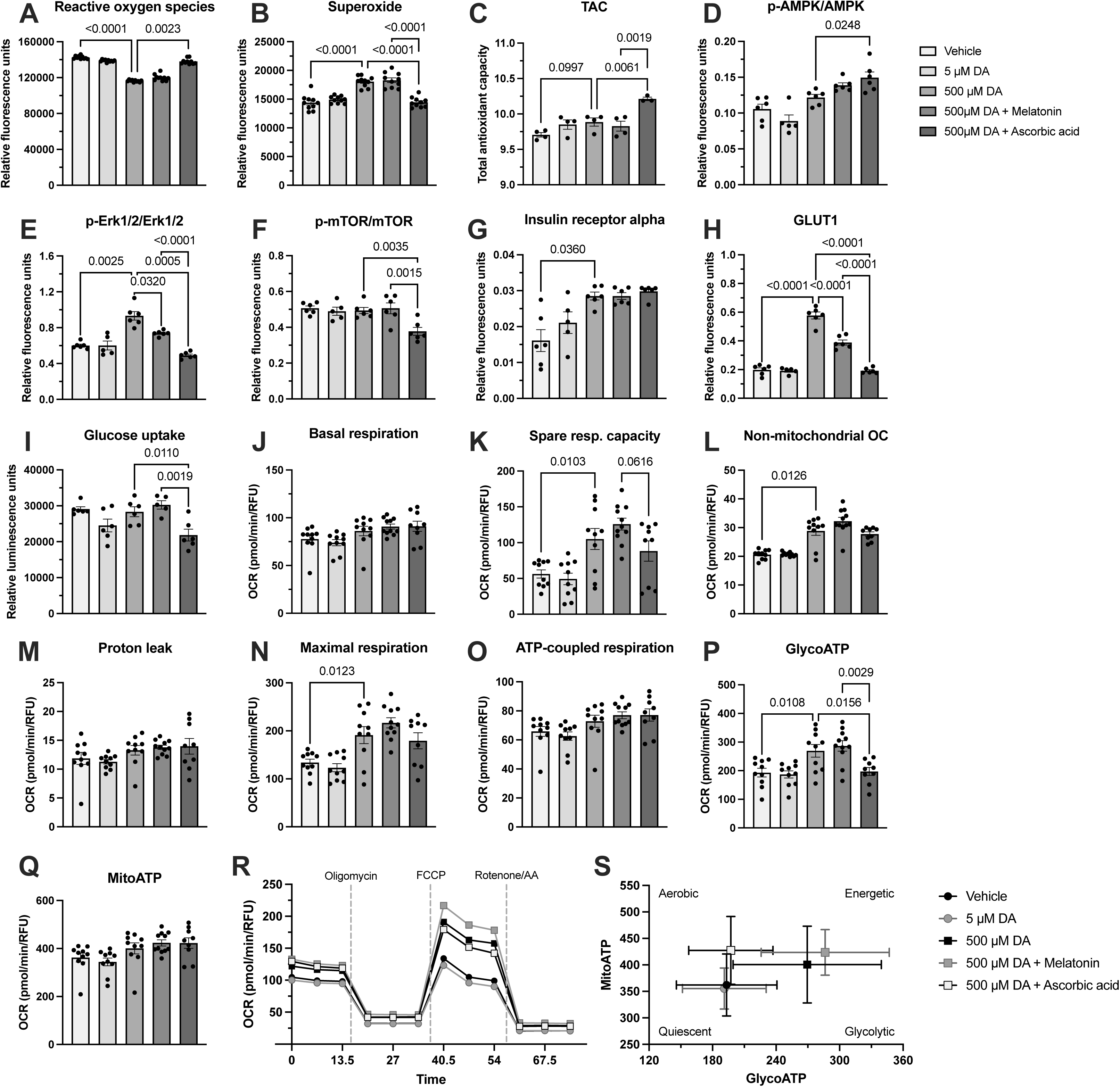
Effects of dopamine (DA), melatonin, and ascorbic acid on oxidative stress and metabolic markers in SH-SY5Y cells. Mean (±SEM) levels (relative fluorescence units (RFU)) of reactive oxygen species *(A)* and superoxide *(B)*; mean (±SEM) total antioxidant capacity (TAC) *(C)*; mean (±SEM) ratios (RFU) of p-AMPK/AMPK *(D)*, p-Erk1/2/Erk1/2 *(E)*, and p-mTOR/mTOR *(F)*; mean (±SEM) levels (RFU) of insulin receptor alpha *(G)* and glucose transporter type 1 (GLUT1) *(H)*; mean (±SEM) levels (relative luminescence units) of glucose uptake *(I)*; mean (±SEM) levels (oxygen consumption rate (OCR) (pmol/min/RFU)) of basal respiration *(J)*, spare respiratory (resp.) capacity *(K)*, non-mitochondrial oxygen consumption (OC) *(L)*, proton leak *(M)*, maximal respiration *(N)*, and ATP-coupled respiration *(O)*; mean (±SEM) levels of basal glycolytic ATP production (glycoATP) (ATP production rate (pmol/min/RFU)) *(P)* and basal mitochondrial ATP production (mitoATP) (ATP production rate) *(Q)* compared between vehicle, 5µM DA, 500µM DA, 500µM DA + melatonin, and 500µM DA + ascorbic acid groups. Effect of treatment is p<0.05 for analyses, as measured by one-way analysis of variance (ANOVA) or Kruskal-Wallis test. Displayed p-values reflect results of Holm-Šídák or Dunn’s for Kruskal-Wallis post-hoc tests. Seahorse analysis kinetic map of OCR vs. time *(R)* and energetic map of MitoATP vs. GlycoATP *(S)*.

### 3.5 500µM dopamine induced a glycolytic high-energy state in SH-SY5Y cells

As dopamine dysregulation has been linked to metabolic reprogramming, we next assessed whether 500µM dopamine drives a glycolytic shift and whether melatonin can prevent any maladaptive changes in mitochondrial function. A significant effect of treatment was found on p-AMPK/AMPK [F (4, 24) = 13.93, p < 0.0001], p-Erk1/2/Erk1/2 [W (4.000, 11.28) = 44.27, p < 0.0001], and p-mTOR/mTOR [F (4, 24) = 6.426, p = 0.0012] ratios, insulin receptor alpha [W (4.000, 11.21) = 5.428, p = 0.0112], and glucose transporter type 1 (GLUT1) [F (4, 24) = 113.7, p < 0.0001] (Figure 3D-H). Post-hoc tests revealed significant increases in p-Erk1/2/Erk1/2, insulin receptor alpha, and GLUT1 in cells treated with 500µM dopamine (Figure 3E, G-H). Pre-treatment with melatonin prior to administration of 500µM dopamine significantly decreased p-Erk1/2/Erk1/2 ratio and GLUT1 when compared to cells treated with 500µM dopamine alone (Figure 3E, H). Cells treated with a combination of ascorbic acid and 500µM dopamine demonstrated significant decreases in p-Erk1/2/Erk1/2 ratio, p-mTOR/mTOR ratio, and GLUT1, and an increase in p-AMPK/AMPK ratio when compared to cells treated with 500µM dopamine alone (Figure 3D-F, H). Significant decreases in p-Erk1/2/Erk1/2, p-mTOR/mTOR, and GLUT1 were seen in cells treated with ascorbic acid and 500µM dopamine when compared to cells treated with melatonin and 500µM dopamine (Figure 3E-F, H). One-way ANOVA analysis also revealed a significant effect of treatment on glucose uptake [F (4, 24) = 6.242, p = 0.0014] (Figure 3I). A significant decrease in glucose uptake was seen in cells treated with ascorbic acid and 500µM dopamine when compared to cells treated with 500µM dopamine alone, and when compared to cells treated with both 500µM dopamine and melatonin (Figure 3I).

One-way ANOVA and Kruskal-Wallis tests revealed a significant effect of treatment on basal respiration [H (4, 45) = 15.68, p = 0.0035], spare respiratory capacity [F (4, 45) = 9.752, p < 0.0001], non-mitochondrial oxygen consumption [H (4, 45) = 32.20, p < 0.0001], proton leak [H (4, 45) = 10.70, p = 0.0302], maximal respiration [F (4, 45) = 10.06, p < 0.0001], ATP-coupled respiration [H (4, 45) = 15.80, p = 0.0033], glycoATP [F (4, 45) = 8.016, p < 0.0001], and mitoATP [H (4, 45) = 15.60, p = 0.0036] (Figure 3J-R). Cells treated with 500µM dopamine demonstrated a significant increase in spare respiratory capacity, non-mitochondrial oxygen consumption, maximal respiration, and glycoATP when compared to cells treated with vehicle (Figure 3K-L, N, P). A significant decrease in glycoATP was seen in cells treated with ascorbic acid and 500µM dopamine when compared to cells treated with 500µM dopamine alone (Figure 3P). Finally, cells treated with 500µM dopamine and ascorbic acid demonstrated significantly reduced glycoATP and a trend reduction in spare respiratory capacity when compared to cells treated with melatonin and 500µM dopamine (Figure 3K, P). Cells treated with 500µM dopamine exhibited a more energetic and glycolytic phenotype than vehicle-treated cells, cells exposed to melatonin prior to 500µM dopamine exhibited a more energetic phenotype than cells treated with 500µM dopamine alone, and cells treated with 500µM dopamine and ascorbic acid exhibited a more aerobic phenotype than cells treated with 500µM dopamine alone (Figure 3S).

One-way ANOVA analyses revealed a significant effect of treatment on DHAP [F (4, 25) = 4.430, p = 0.0076], fructose 1,6-bisphosphate (F16DP) [F (4, 25) = 2.840, p = 0.0454], fructose 6-phosphate (F6P) [F (4, 25) = 3.619, p = 0.0185], GA3P [F (4, 25) = 4.721, p = 0.0056], glycerol 3-phosphate (G3P) [W (4.000, 12.42) = 13.47, p = 0.0002], D-ribose 5-phosphate (R5P) [F(4, 25) = 3.254, p = 0.0280], D-ribulose 5-phosphate (RL5P) [F (4, 25) = 2.976, p = 0.0387], sedoheptulose 7-phosphate (SH7P) [F(4, 25) = 2.858, p = 0.0444], coenzyme A (CoA) [F (4, 25) = 3.300, p = 0.0266], creatine phosphate (CrePO4) [F (4, 25) = 8.965, p = 0.0001], oxidised nicotinamide adenine dinucleotide (NAD) [F (4, 25) = 2.993, p = 0.0379], and uridine diphosphate N-acetylglucosamine (UDPNAG) [F (4, 25) = 6.084, p = 0.0015]; and a trend effect on 3-phospho-D-glycerate (3PG) [F (4, 25) = 2.705, p = 0.0533] and D-xylulose 5-phosphate (X5P) [F (4, 25) = 2.189, p = 0.0994] (Figure 4A-N). Cells treated with 5µM dopamine demonstrated a decrease in NAD when compared to vehicle-treated cells (Figure 4M). Cells treated with 500µM dopamine demonstrated significant decreases in G3P and CoA, and a trend decrease in SH7P when compared to vehicle-treated cells (Figure 4F, I, K). When compared to cells treated with 500µM dopamine alone, cells treated with 500µM dopamine and ascorbic acid demonstrated significant increases in DHAP, F16DP, GA3P, G3P, and R5P, and significant decreases in CrePO4 and UDPNAG (Figure 4B-C, E-G, L, N). Finally, significant increases in DHAP and R5P, a trend increase in GA3P, and significant decreases in UDPNAG and CrePO4 were seen in cells treated with ascorbic acid and 500µM dopamine when compared to cells treated with melatonin and 500µM dopamine (Figure 4B, E, G, N, L). These findings indicate that excess dopamine generates a metabolically activated, glycolytic phenotype, and that melatonin only partially protects against dopamine-induced metabolic reprogramming.

**Figure 4:**
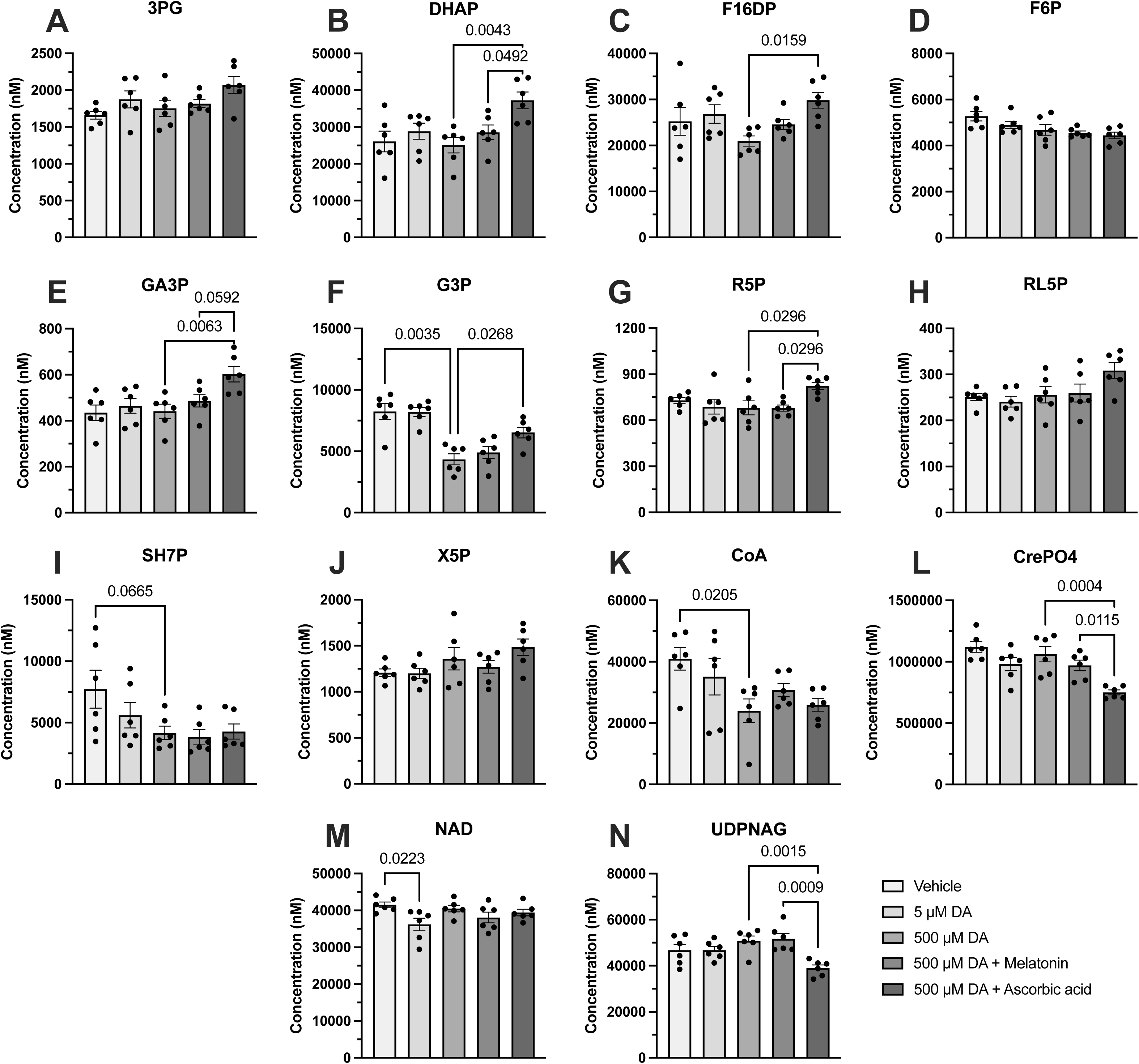
Effects of dopamine (DA), melatonin, and ascorbic acid on central carbon metabolites (CCM) in SH-SY5Y cells. Mean (±SEM) concentration (nM) of 3-phospho-D-glycerate (3PG) ***(A)***, dihydroxyacetone phosphate (DHAP) ***(B)***, fructose 1,6-bisphosphate (F16DP) ***(C)*** fructose 6-phosphate (F6P) ***(D)***, glyceraldehyde 3-phosphate (GA3P) ***(E)***, glycerol 3-phosphate (G3P) ***(F)***, D-ribose 5-phosphate (R5P) ***(G)***, D-ribulose 5-phosphate (RL5P) ***(H)***, sedoheptulose 7-phosphate (SH7P) ***(I)*** D-xylulose 5-phosphate (X5P) ***(J)***, coenzyme A (CoA) ***(K)***, creatine phosphate (CrePO4) ***(L)***, oxidised nicotinamide adenine dinucleotide (NAD) ***(M)***, and uridine diphosphate N-acetylglucosamine (UDPNAG) ***(N)*** compared between vehicle, 5µM DA, 500µM DA, 500µM DA + melatonin, and 500µM DA + ascorbic acid groups. Effect of treatment is p<0.1 for analyses, as measured by one-way analysis of variance (ANOVA) or Welch’s ANOVA. Displayed p-values reflect results of Holm-Šídák or Dunnett’s T3 post-hoc tests.

## 4 Discussion

In this study, we investigated whether melatonin is capable of mitigating metabolic disturbance, oxidative stress, and dopaminergic dysfunction in neuronal models of acute inflammation and hyperdopaminergia. While the main focus of this study was to improve our understanding of the therapeutic mechanisms of melatonin in BD, the results also have important implications for other conditions where hypomelatoninaemia, inflammation, and dopamine dysregulation play an important role, such as PD and Alzheimer’s disease (Martorana & Koch, 2014; Ribeiro et al., 2025).

The results of this study demonstrate that melatonin can mitigate IL-6 and dopamine-induced oxidative stress and metabolic dysfunction in differentiated SH-SY5Y cells. These findings support the role of melatonin as a regulator of redox balance and cellular energy homeostasis, with implications for understanding the molecular underpinnings of BD and related neuropsychiatric conditions. Our results suggest that IL-6, a pro-inflammatory cytokine that is elevated across all mood states in BD, may induce a hyperdopaminergic effect in neuronal cells, as evidenced by increased intracellular dopamine content and decreased dopamine turnover. This contrasts with prior evidence showing that inflammation generally suppresses dopamine signalling (Felger & Treadway, 2017), suggesting that IL-6’s effects may be context-dependent, potentially influenced by exposure duration or cell maturity. Notably, melatonin pretreatment suppressed IL-6-induced dopamine accumulation, consistent with melatonin’s proposed antidopaminergic and immunomodulatory role (Chitimus et al., 2020; Pe et al., 2025).

Melatonin also mitigated IL-6-induced increases in AMPK signalling, restored mitochondrial capacity, and reduced non-mitochondrial oxygen consumption, findings aligned with its known capacity to regulate metabolism and improve mitochondrial function (Cardinali & Vigo, 2017; Jiki et al., 2018). These metabolic benefits were accompanied by enhanced glucose uptake and a shift toward a quiescent energetic profile, indicating improved metabolic efficiency under inflammatory stress. Melatonin’s ability to attenuate dopamine-induced metabolic stress via Erk1/2 signalling and GLUT1 further supports its neuroprotective profile. However, antioxidant control experiments with ascorbic acid suggest melatonin’s effects extend beyond ROS scavenging. Although ascorbic acid had positive effects on select oxidative and metabolic stress markers, it also increased p-AMPK/AMPK, reduced glucose uptake, reduced spare respiratory capacity, and increased glycolytic intermediates, supporting an overall negative effect on mitochondrial health and function (Pe et al., 2025).

Collectively, our results support a model in which melatonin acts through converging mechanisms, including redox modulation, metabolic reprogramming, and dopaminergic-inflammation crosstalk, to protect neurons from stress-related dysfunction. These findings position melatonin as a potential therapeutic adjunct in disorders marked by inflammation, oxidative stress, and monoamine imbalance, such as BD (Ribeiro et al., 2025).

### 4.1 Melatonin suppresses inflammation-induced hyperdopaminergia

Impairments in the crosstalk between the immune system and dopamine – known as the dopamine-immune axis – are believed to contribute to a number of psychiatric disorders, including BD (Felger & Treadway, 2017; Treadway, Cooper, & Miller, 2019). Preclinical and clinical research both support a critical pathophysiological role of chronic inflammation in disrupting dopamine neurotransmission via alterations in dopamine synthesis, release, and reuptake mechanisms (Treadway et al., 2019). Cytokines such as IL-6 are believed to be essential for the propagation of peripheral inflammatory signals to the brain, where they directly act upon dopamine neurons or alter dopamine neuron function via activation of neuroimmune cells (e.g., microglia) (Treadway et al., 2019; Varela et al., 2025). Chronic administration of IL-6 has been shown to disrupt dopamine synthesis and release in *ex vivo* and *in vitro* models (Emmons, Wallace, & Fordahl, 2023; R. Li et al., 2012; W. Li, Knowlton, Woodward, & Habecker, 2003). Furthermore, peripheral IL-6 levels correlated with decreased functional corticostriatal connectivity in depressed patients, suggesting a negative association between IL-6 and dopaminergic neurotransmission (Felger et al., 2016).

Due to this evidence, we hypothesised that IL-6 treatment would significantly reduce TH and/or dopamine content in SH-SY5Y cells. Instead, IL-6 significantly increased dopamine content without affecting TH. Acute administration of IL-6 has been shown to augment hippocampal and prefrontal dopamine utilisation, TH activity (i.e., the capacity to convert tyrosine to dopamine) in adrenal medullary chromaffin cells, and dopamine-dependent behaviours in mice, demonstrating that IL-6 can enhance dopamine function (Jenkins et al., 2016; Zalcman et al., 1994; Zalcman, Murray, Dyck, Greenberg, & Nance, 1998). Furthermore, investigations into the effects of IL-6 on dopaminergic markers have not utilised SH-SY5Y cells. Therefore, the results of this study suggest that subchronic IL-6 administration may have a hyperdopaminergic effect that is specific to adult dopaminergic neurons. This novel mechanism may help to explain why acute inflammation can trigger BD in patients with no past psychiatric history. For instance, severe SARS-CoV-2 infection, which is associated with significant increases in IL-6, has been shown to precipitate first-time manic episodes in numerous case studies (Costa, Roman Meller, & Kapczinski, 2023; D’Imperio, Lo, Bettini, Prada, & Bondolfi, 2022; Majidpoor & Mortezaee, 2022; Sprenger, Bare, Kashyap, & Cardella, 2022).

Additionally, IL-6 decreased dopamine turnover, as indicated by reduced HVA/dopamine ratio. Impaired degradation of dopamine may therefore underpin the hyperdopaminergic effect of IL-6 on SH-SY5Y cells. A reduction in dopamine turnover has also been seen in the striatum of patients treated with interferon-alpha, as well as postmortem occipital cortex samples obtained from patients with BD (Capuron et al., 2012; Young, Warsh, Kish, Shannak, & Hornykeiwicz, 1994). IL-6 signalling has been shown to downregulate monoamine oxidase A, the primary degradation enzyme of dopamine (Huang et al., 2012); however, no changes in MAOA gene expression were seen in this study. A trend decrease in SV2C, which is a marker of dopamine release, was also seen with IL-6 treatment. The reduction in SV2C seen in this study could be a result of negative feedback, or may indicate a pattern of dopamine dysregulation whereby metabolism and release are impaired, leading to a build-up of intracellular dopamine.

We also hypothesised that melatonin treatment would reduce dopaminergic markers in this study, due to its documented antagonistic effect on dopamine. Melatonin has been shown to significantly reduce dopamine release in *ex vivo* rodent striatum, brainstem, cerebral cortex, cerebellum, hippocampus, and hypothalamus slice cultures (Zisapel, Egozi, & Laudon, 1982, 1983, 1985; Zisapel & Laudon, 1982, 1983, 1987). Preclinical research also supports a significant inhibitory effect of melatonin on dopamine synthesis (Alexiuk & Vriend, 1991, 2007; Shieh, Chu, & Pan, 1997). There were no significant effects of melatonin on dopaminergic markers in this study; however, pretreatment with melatonin prior to IL-6 inhibited the increase in dopamine content seen in cells treated with IL-6 alone. This result complements findings from preclinical studies, which have demonstrated suppression of stress-, amphetamine-, nicotine-, and cocaine-evoked increases in dopamine content, release, or synthesis by melatonin and agomelatine, a non-selective melatonin receptor agonist (Alexiuk & Vriend, 1991, 2007; Barbosa-Mendez, Perez-Sanchez, & Salazar-Juarez, 2023; Exposito, Mora, & Oaknin, 1995; Exposito, Mora, Zisapel, & Oaknin, 1995; Park, Jung, Park, Yang, & Kim, 2018; Schiller, Champney, Reiter, & Dohrman, 2003). Overall, the results of this study show, for the first time, that melatonin can reduce hyperdopaminergia induced by an inflammatory stimulus in a dopaminergic neuronal model. Interestingly, like IL-6, melatonin also reduced dopamine turnover. However, the effect of melatonin on dopamine turnover may be related to its role as an antioxidant, as the conversion of dopamine to HVA generates ROS (Meiser, Weindl, & Hiller, 2013).

### 4.2 IL-6 induces metabolic stress and mitochondrial dysfunction

Altered immunometabolism – the link between the immune system and metabolic processes – has been shown to play a role in the pathophysiology of mood disorders, including BD (Teixeira, Scholl, & Bauer, 2025). Chronic inflammation significantly increases energy requirements, leading to metabolic reprogramming, whereby metabolic pathways shift from oxidative phosphorylation to glycolysis in a manner similar to the Warburg effect seen in cancer cells (Bernier, York, & MacVicar, 2020; Garaude, 2019; Teixeira et al., 2025). Metabolic reprogramming is also believed to play a role in mitochondrial dysregulation in BD, and is therefore believed to be central to its aetiology and pathophysiology (Morris et al., 2017).

IL-6 has been shown to alter many metabolic processes, and systemic, excessive IL-6 has been linked to insulin resistance, glucose intolerance, hyperglycaemia, dyslipidaemia, and central obesity (Y. S. Li, Ren, & Cao, 2022). In this study, IL-6 increased p-AMPK/AMPK and p-Erk1/2/Erk1/2 ratios, both of which are markers of impaired glucose uptake, increased glycolysis, mitochondrial dysfunction and metabolic stress (Martin-Vega & Cobb, 2025; Mihaylova & Shaw, 2011; Toyama et al., 2016). IL-6 also significantly depressed spare respiratory capacity, an indicator of mitochondrial fitness and plasticity (Marchetti, Fovez, Germain, Khamari, & Kluza, 2020), and increased TAC, which indicates an adaptive response to oxidative stress (Silvestrini & Mancini, 2024). Overall, these findings show that IL-6 induces metabolic and oxidative stress, and impairs mitochondrial function and glucose homeostasis in differentiated dopaminergic neurons.

Furthermore, IL-6 was shown to significantly increase nucleotide mono- and diphosphates, without affecting nucleotide triphosphates. This may indicate increases in nucleotide triphosphate turnover, as is seen in high-energy states (such as aerobic exercise) (Alghannam, Ghaith, & Alhussain, 2021). However, increases in the AMP/ATP ratio, particularly when accompanied by increases in AMPK activation, can also indicate metabolic stress (Lin & Hardie, 2018). IL-6 was also shown to decrease acetyl CoA, which can occur as a result of AMPK activation during periods of depleted cellular nutrient availability (Cholico et al., 2023). These results suggest that IL-6 induces a dysfunctional hyperenergetic state in neurons with subsequent exhaustion, similar to what is seen in BD. Therefore, this study may have important implications for understanding the role of IL-6 in modulating neurometabolism in psychiatric disease.

### 4.3 Melatonin reduces metabolic and oxidative stress and improves mitochondrial function

Melatonin is known to play an important role in the regulation of energy metabolism, and has widespread effects on insulin signalling, energy expenditure and storage, glucose uptake, and lipid metabolism (Cipolla-Neto, Amaral, Afeche, Tan, & Reiter, 2014). The reduction in melatonin production during ageing and shift-work is thought to augment insulin resistance, glucose intolerance, dyslipidaemia, and energy imbalance, and ultimately lead to metabolic diseases such as type 2 diabetes and obesity (Cipolla-Neto et al., 2014). Hypomelatoninaemia may therefore be a key contributor towards the systemic metabolic disturbances seen in BD (Cipolla-Neto et al., 2014; Morris et al., 2017).

Treatment with melatonin alone decreased the p-AMPK/AMPK ratio in differentiated SH-SY5Y cells, indicating a reduction in metabolic stress. Melatonin also reduced G1P, a key intermediate in the breakdown of glycogen to glucose, suggesting an increase in glycogenesis and/or a decrease in glycogenolysis and gluconeogenesis (Dashty, 2013). Increased glycogenolysis and decreased glycogenesis contribute to hyperglycaemia and are markers of insulin resistance (Barroso, Jurado-Aguilar, Wahli, Palomer, & Vazquez-Carrera, 2024). Like IL-6, melatonin increased nucleotide mono- and diphosphates without affecting nucleotide triphosphates, which suggests increased nucleotide triphosphate turnover. As p-AMPK/AMPK was also reduced, our findings indicate that melatonin induces a high-energy state utilising more effective mechanisms for ATP production, unrelated to metabolic stress and maladaptive reprogramming. Melatonin treatment was also shown to reduce ROS and increase NADH, a directly-acting antioxidant (Kirsch & De Groot, 2001), thus demonstrating a reduction in oxidative stress.

Melatonin also appears to have a protective role against IL-6-induced mitochondrial and metabolic dysfunction, as illustrated by a reduction in p-AMPK/AMPK ratio and increase in spare respiratory capacity, glucose uptake, and NADH when compared to cells treated with IL-6 only. Furthermore, melatonin pretreatment reduced non-mitochondrial oxygen consumption, which is considered to be a marker of poor bioenergetic health and metabolic/oxidative stress (Chacko et al., 2014). A reduction in TAC was seen in cells treated with melatonin and IL-6, which can indicate high antioxidant consumption in response to oxidative stress (Silvestrini, Meucci, Ricerca, & Mancini, 2023). Melatonin pretreatment also activated GSK3β (as demonstrated by reduced p-GSK3β/GSK3β ratio); this result complements prior research (Ali & Kim, 2015; Fu et al., 2022). Increased GSK3β signalling has been implicated in the development of BD (Ali & Kim, 2015; Won & Kim, 2017); however, activation of GSK3β has also been shown to reduce glycolysis within the brain via regulation of hexokinase 2 (Wang, Li, & Di, 2022). GSK3β also promotes and stabilises a number of circadian proteins linked to melatonin function (Abreu & Braganca, 2015). Increases in GA3P and DHAP, two interconverting molecules involved in fructose metabolism and glycolysis, were also seen in cells exposed to melatonin prior to IL-6 (Dashty, 2013). Melatonin has been shown to increase DHAP in a cellular model of peritoneal fibrosis; it was theorised that this increase was pivotal to cellular survival (Ruan et al., 2024). Altogether, the results of this study suggest that melatonin improves mitochondrial function and reduces metabolic and oxidative stress in IL-6-treated differentiated SH-SY5Y cells.

### 4.4 Excess dopamine induces a glycolytic high-energy state

As IL-6 increased dopamine content and induced mitochondrial dysfunction in SH-SY5Y cells, we postulated that the metabolic effects of IL-6 may have arisen from hyperdopaminergia. Therefore, we examined the effects of high dopamine on metabolism and oxidative stress in differentiated SH-SY5Y cells. Furthermore, as melatonin reduced metabolic and oxidative stress, and improved mitochondrial function in IL-6-treated cells, it was theorised that melatonin may have similar effects in dopamine-treated adult dopaminergic neurons. Finally, the effects of melatonin were compared to ascorbic acid in dopamine-treated cells, to determine whether the effects of dopamine and melatonin on metabolism could be solely attributed to oxidative stress and antioxidation.

In this study, high dose (500µM) dopamine increased p-Erk1/2/Erk1/2, insulin receptor alpha and GLUT1 expression. Overexpression of both p-Erk1/2/Erk1/2 and GLUT1 are indicative of increased glycolysis (Massari et al., 2016; Papa, Choy, & Bubici, 2019). Increases in insulin receptor alpha typically indicate insulin resistance, a common marker of BD (Cuperfain, Kennedy, & Goncalves, 2020; Haeusler, McGraw, & Accili, 2018); however, insulin signalling has also been shown to play a crucial role in the metabolic reprogramming of immune cells towards glycolysis (van Niekerk, Christowitz, Conradie, & Engelbrecht, 2020). High dose dopamine also decreased levels of CoA, a key metabolic cofactor which has been shown to facilitate antioxidation in response to oxidative or metabolic stress (Filonenko & Gout, 2023); and G3P, a key intermediate in lipid metabolism, which may indicate a shift away from lipid synthesis and towards glycolysis (Mracek, Drahota, & Houstek, 2013). Furthermore, high dose dopamine increased glycoATP, maximal respiration, and non-mitochondrial oxygen consumption. Interestingly, spare respiratory capacity was increased in cells treated with high levels of dopamine, which distinguishes the metabolic phenotype of dopamine-treated neurons from cancer cells, and suggest that twelve hours of dopamine may induce metabolic programming without compromising mitochondrial health (Marchetti et al., 2020). Overall, these results indicate that excess dopamine may induce a predominantly glycolytic, dysregulated, high-energy state in adult dopaminergic neurons, in a manner similar to mania (Morris et al., 2017). This high-energy state would be unsustainable and detrimental in the long-term, which may lead to a bioenergetic deficit and consequent mood shift. Therefore, this study may provide novel insight into the connection between hyperdopaminergia and high-energy states in psychiatric disease, with particular relevance to BD.

As dopamine catabolism is known to generate ROS and SOX, it was hypothesised that excess dopamine would increase ROS and SOX levels in differentiated SH-SY5Y cells. High dose dopamine increased SOX levels, as expected, as enzymatic oxidation of dopamine generates SOX radical anions (Meiser, Weindl, & Hiller, 2013). However, ROS levels decreased in 500µM dopamine-treated cells. This could be due to a number of factors. For instance, dopamine degradation may have increased the levels of other non-ROS substances that contribute to oxidative stress instead of ROS, such as quinones, which were not detectable using the ROS/SOX assay (Meiser, Weindl, & Hiller, 2013). Furthermore, TAC was increased at a trend level, which may suggest that application of dopamine leads to adaptive augmentation of non-enzymatic antioxidants as a method of combatting oxidative stress. In order to further our understanding of the effects of excess dopamine on oxidative stress, it may be beneficial to repeat this experiment with other oxidative stress measures, such as malondialdehyde, protein carbonyl content, or DNA damage (Azzi, 2022).

### 4.5 Melatonin partially mitigates the metabolic effects of dopamine while ascorbic acid does not

The final aim of this study was to examine the individual effects of melatonin and ascorbic acid on metabolism and oxidative stress in dopamine-treated SH-SY5Y cells, and compare the two compounds to determine whether antioxidation underlies the protective effects of melatonin. Treatment with melatonin prior to excess dopamine was shown to reduce p-Erk1/2/Erk1/2 and GLUT1, demonstrating a modest role of melatonin in reversing metabolic reprogramming in dopamine-treated SH-SY5Y cells. Melatonin was not shown to have any other effects in this cell model, which suggest that melatonin’s effects are more selective and impact more general processes rather than dopamine-specific mechanisms of action. These outcomes further illustrate that IL-6 and dopamine affect metabolism and mitochondrial function through differing pathways. Melatonin may therefore have a mitigating effect on manic symptomology in BD patients directly by reducing dopamine, and indirectly by moderating the metabolic effects of inflammation.

It was theorised that ascorbic acid would reverse any deleterious effects of dopamine on metabolism and oxidative stress due to its potent antioxidant properties, and that melatonin and ascorbic acid would have similar effects. Ascorbic acid was shown to increase G3P and reduce p-Erk1/2/Erk1/2, p-mTOR/mTOR, GLUT1, and glycoATP in dopamine-treated cells, which altogether support a role of ascorbic acid in reducing glycolysis (Saxton & Sabatini, 2017). Additionally, ascorbic acid may have reduced hexosamine biosynthesis pathway (HBP) activity, as indicated by lowered UDPNAG (Paneque, Fortus, Zheng, Werlen, & Jacinto, 2023). Increased HBP is associated with insulin resistance and cancer (Paneque et al., 2023). However, ascorbic acid also significantly increased p-AMPK/AMPK and reduced glucose uptake, which suggest that ascorbic acid increases metabolic stress in dopamine-treated cells. Ascorbic acid also increased GA3P, DHAP, 3PG and F16DP, all of which are glycolytic intermediates, which provides contradictory evidence that ascorbic acid increases glycolysis (Dashty, 2013). Furthermore, ascorbic acid reduced CrePO4, which plays a crucial role in replenishing ATP in times of high energy utilisation (Gaddi, Galuppo, & Yang, 2017). Ascorbic acid may also increase glucose flux through the pentose phosphate pathway (PPP), as illustrated by increases in the PPP intermediates R5P and R5P (Dashty, 2013). Other studies have found a similar effect of ascorbic acid on the PPP, which is activated during periods of oxidative stress, as the final product NADPH provides antioxidant defence through the reduction of oxidised glutathione (Williams & Ford, 2004).

Similarly, ascorbic acid significantly reduced SOX, suggesting a suppressive effect on oxidative stress; however, ROS levels were significantly increased when compared to dopamine-treated cells. This may again reflect the limitations of the ROS/SOX assay and emphasises the need for further testing with other measurements of oxidative stress. High dose ascorbic acid has been theorised to increase oxidative stress (Williams & Ford, 2004; Zhang et al., 2016); however, the dose chosen in this study was lower than what has previously been shown to have antioxidant properties in SH-SY5Y cells (Gomez-Santos et al., 2003; Lai & Yu, 1997). Overall, it is clear that melatonin and ascorbic acid had divergent effects on metabolism in differentiated SH-SY5Y cells treated with high-dose dopamine. This demonstrates that the effects of melatonin on metabolism cannot be attributed to its antioxidant effects, and that dopamine affects mitochondrial function via mechanisms independent of oxidative stress.

### 4.6 Limitations and future directions

There are several limitations that need to be taken into consideration when interpreting experimental results. First, while SH-SY5Y cells are regularly used in neurobiological research, particularly as a model of PD, there are known disadvantages associated with the cell line, including its origin as a cancer and that it is not considered to be “purely dopaminergic” (Xicoy, Wieringa, & Martens, 2017). Secondly, dopamine intake and release mechanisms were not measured in this study. Further analysis of dopaminergic mechanisms would aid in the understanding of these results. Similarly, the ROS/SOX assay and TAC results are difficult to interpret by themselves, and further investigations into the effects of IL-6, dopamine, and melatonin on other measures of oxidative stress would be beneficial. Finally, the effects of ascorbic acid were not tested in cells treated with IL-6.

The results of this study provide many new and interesting future avenues of research, and highlight the potential for melatonin as a treatment for BD. Investigation of the effects of IL-6, melatonin, and dopamine on other aspects of metabolism and mitochondrial health in dopaminergic neurons, such as morphological analysis of mitochondria and quantification of electron transport chain complexes, would further elucidate the cellular mechanisms connecting inflammation, dopamine dysregulation, melatonin, and metabolic reprogramming. Furthermore, it would be beneficial to evaluate other measurements of oxidative stress, such as quinones, protein carbonyl content, reactive nitrogen species, lipid peroxidation markers, and DNA damage in the same cellular models. Finally, translating these experiments to animal models of excess dopamine and IL-6 is the essential next step in understanding the role of melatonin in treating neurodegenerative and psychiatric diseases.

## 5 Conclusion

This study provides new evidence that IL-6 causes dopamine dysfunction, metabolic stress, and mitochondrial impairment in differentiated SH-SY5Y cells, and that melatonin has the capacity to prevent IL-6-induced hyperdopaminergia, oxidative stress, and metabolic reprogramming. Furthermore, this study demonstrated that excess dopamine induces a glycolytic high-energy state in dopaminergic neurons, thereby providing a crucial connection between hyperdopaminergia and dysregulated bioenergetics in mania. Finally, it was shown that melatonin has a partial protective effect against the effects of dopamine on metabolic reprogramming, but treatment with a potent antioxidant (ascorbic acid) significantly worsened metabolic stress and mitochondrial dysfunction in cells treated with excess dopamine. Thus, the effects of melatonin on metabolism cannot be attributed to its antioxidant potential alone. Together, these findings illustrate the important links between inflammation, dopamine dysregulation and metabolic dysfunction in psychiatric disease, and provide new evidence as to the mechanisms underpinning melatonin’s efficacy as a therapeutic for BD.

## Acknowledgements

The authors would like to acknowledge the support and assistance provided by the University of Queensland Metabolomics and Proteomics (Q-Map) team for the completion of this study.

---

**Supplementary Data 1:** Details of reagents, kits, software, equipment, and instruments.

| Item | Brand | Country | Catalogue number |
| --- | --- | --- | --- |
| **Reagents** |  |  |  |
| SH-SY5Y cell line | Sigma-Aldrich, Merck | Germany | #94030304 |
| Dulbecco's Modified Eagle Medium/Nutrient Mixture F-12 (DMEM/F12) | Gibco, Thermo Fisher Scientific Inc. | United States | #11320033 |
| Heat-inactivated foetal bovine serum (FBS) | Gibco, Thermo Fisher Scientific Inc. | United States | #10099141 |
| GlutaMAX™ 100X | Gibco, Thermo Fisher Scientific Inc. | United States | #35050061 |
| 10000 units/ml penicillin-streptomycin | Gibco, Thermo Fisher Scientific Inc. | United States | #15140122 |
| Poly-D-lysine hydrobromide | Sigma-Aldrich, Merck | Germany | #P6407-5MG |
| B27™ Supplement (50X), serum free | Gibco, Thermo Fisher Scientific Inc. | United States | #17504044 |
| All-trans retinoic acid (RA) | Sigma-Aldrich, Merck | Germany | #R2625 |
| Melatonin | Sigma-Aldrich, Merck | Germany | #M5250-1G |
| Interleukin-6 (IL-6) human | Sigma-Aldrich, Merck | Germany | #I1395 |
| Dopamine | Sigma-Aldrich, Merck | Germany | #H8502-5G |
| L-ascorbic acid | Supelco, Merck | Germany | #PHR1008 |
| Intercept (PBS) Blocking Buffer | Li-Cor | United States | #927-70001 |
| IRDye 800CW Goat anti-rabbit IgG secondary antibody | Li-Cor | United States | #925-32211 |
| Cell Tag 700 Stain | Li-Cor | United States | #926-41090 |
| Seahorse XF DMEM, pH 7.4 | Agilent Technologies Inc. | United States | #103575-100 |
| Seahorse XF 200mM glutamine | Agilent Technologies Inc. | United States | #103579-100 |
| 100mM sodium pyruvate | Thermo Fisher Scientific Inc. | United States | #11360-070 |
| 200g/L D-glucose | Thermo Fisher Scientific Inc. | United States | #A24940-01 |
| Trypsin-EDTA (0.25%), phenol red | Gibco, Thermo Fisher Scientific Inc. | United States | #25200056 |
| **Kits** |  |  |  |
| RNeasy® Mini kit | QIAGEN | Germany | #74106 |
| RNAse-free DNase kit | QIAGEN | Germany | #79256 |
| SensiFAST™ cDNA synthesis kit | Bioline, Meridian Bioscience | United States | #BIO-65054 |
| SensiFAST™ SYBR® No-ROX kit | Bioline, Meridian Bioscience | United States | #BIO-98020 |
| Glucose Uptake-Glo™ Assay | Promega | United States | #J1342 |
| Seahorse XF Cell Mito Stress Test assay | Agilent Technologies Inc. | United States | #103015-100 |
| ROS/Superoxide Detection Assay kit (Cell-based) | abcam | United Kingdom | #ab139476 |
| Total Antioxidant Capacity Assay Kit | abcam | United Kingdom | #ab65329 |
| ***Instruments*** |  |  |  |
| NanoPhotometer N60 microvolume spectrophotometer | Implen GmbH | Germany | N/A |
| LightCycler<sup>®</sup> 480 Instrument II | Roche Diagnostics | Switzerland | #05015278001 |
| Coulochem III electrochemical detector | Thermo Fisher Scientific Inc. | United States | N/A |
| Model 5014B Microdialysis High Performance Analytical Cell | ESA, Thermo Fisher Scientific Inc. | United States | #70-0520B |
| Odyssey CLx Imaging system | Li-Cor | United States | N/A |
| Nexera ultra high-performance liquid chromatograph | Shimadzu | Japan | N/A |
| 8060 triple quadrupole mass spectrometer | Shimadzu | Japan | N/A |
| CLARIOstar Plus microplate reader | BMG LABTECH GmbH | Germany | N/A |
| Seahorse XFp Analyzer | Agilent Technologies Inc. | United States | #S7802A |
| ***Equipment*** |  |  |  |
| 96-well Seahorse XFe96 Pro Cell Culture Microplate | Agilent Technologies Inc. | United States | #103794-100 |
| SunFire C8 3µm, 4.6 x 100 mm columns | Waters Corporation | United States | N/A |
| Acquity HSS T3 1.8µm, 2.1mm x 100mm columns | Waters Corporation | United States | #186003539 |
| Acquity HSS T3 1.8µm 2.1mm x 5mm guard columns | Waters Corporation | United States | #186003976 |
| ***Software*** |  |  |  |
| Chemstation 7.2 | Agilent Technologies Inc. | United States | N/A |
| Agilent Seahorse Analytics | Agilent Technologies Inc. | United States | N/A |
| GraphPad Prism 10.0.2 | GraphPad Software Inc. | United States | N/A |

**Supplementary Data 2:**
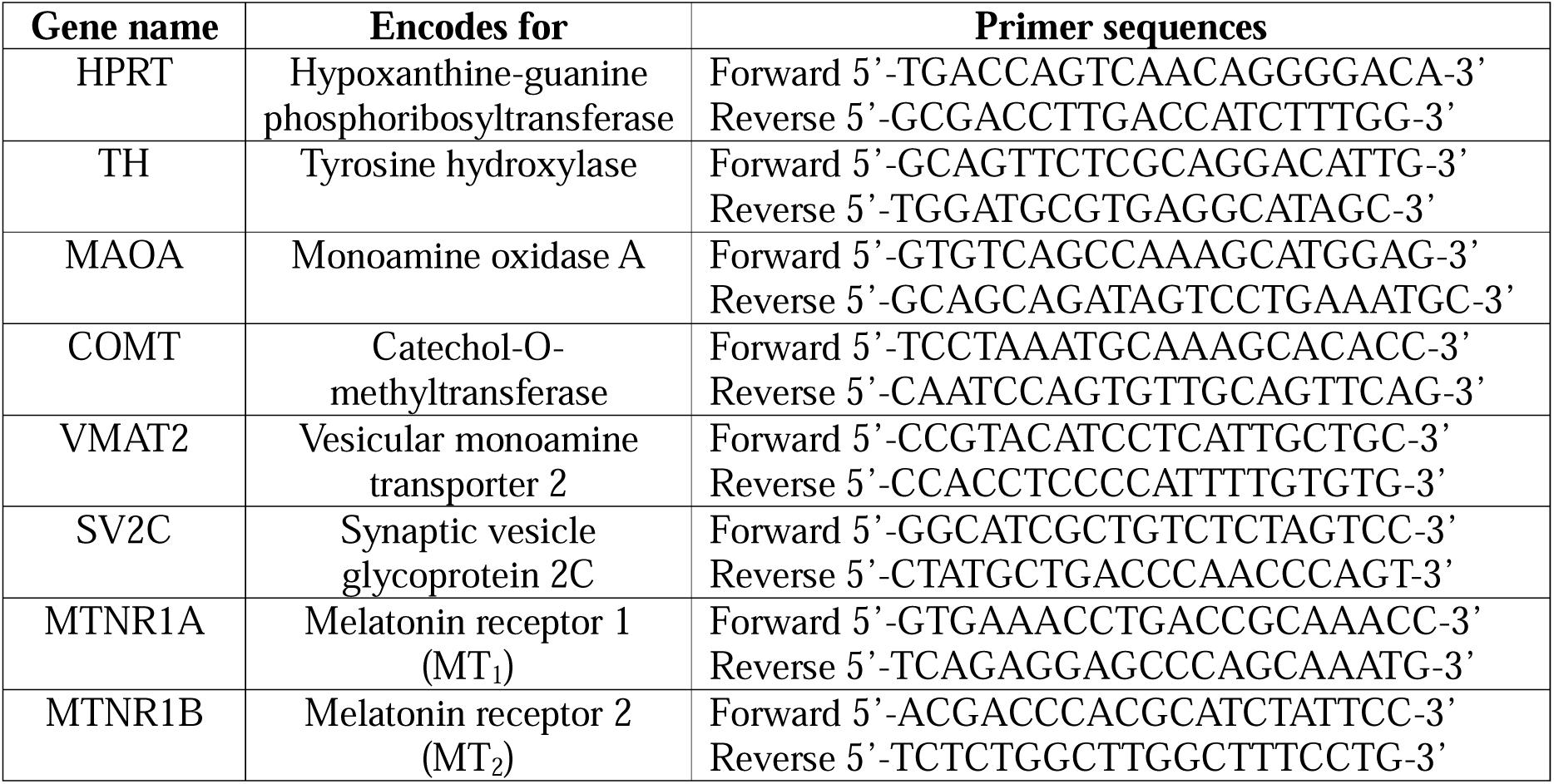
Primer details for quantitative real-time PCR.

**Supplementary Data 3:** Primary antibodies used in In-cell Western assays.

| **Antibody** | **Dilution** | **Species** | **Source** | **Category number** |
| --- | --- | --- | --- | --- |
| Insulin receptor alpha (IRα) | 1:150 | Rabbit | Cell Signalling Technology, United States | #74118 |
| Glucose transporter 1 (GLUT1) | 1:1000 | Rabbit | Abcam, United Kingdom | #ab115730 |
| p44/42 MAPK (Erk1/2) | 1:200 | Rabbit | Cell Signalling Technology, United States | #4695 |
| Phospho-p44/42 MAPK (p-Erk1/2) | 1:200 | Rabbit | Cell Signalling Technology, United States | #4376 |
| Glycogen synthase kinase-3 beta (GSK-3β) | 1:200 | Rabbit | Cell Signalling Technology, United States | #8213 |
| Phospho-glycogen synthase-3 beta (p-GSK-3β) | 1:200 | Rabbit | Cell Signalling Technology, United States | #8213 |
| AMP-activated protein kinase alpha (AMPKα) | 1:200 | Rabbit | Cell Signalling Technology, United States | #2532 |
| Phospho-AMP-activated protein kinase alpha (p-AMPKα) | 1:200 | Rabbit | Cell Signalling Technology, United States | #2535 |
| Mammalian target of rapamycin (mTOR) | 1:200 | Rabbit | Thermo Fisher Scientific Inc., United States | #PA5-34663 |
| Phospho-mammalian target of rapamycin (p-mTOR) | 1:200 | Rabbit | Cell Signalling Technology, United States | #5536 |

**Supplementary Data 4:** Central carbon metabolism (CCM) compound list.

| **Abbreviation** | **Analytical compound** | **Molecular Formula** |
| --- | --- | --- |
| 2PG | 2-phospho-D-glyceric acid | C<sub>3</sub>H<sub>7</sub>O<sub>7</sub>P |
| 3PG | 3- phospho-D-glyceric acid | C<sub>3</sub>H<sub>7</sub>O<sub>7</sub>P |
| 3DHQ | 3-dehydroquinic acid | C<sub>7</sub>H<sub>10</sub>O<sub>6</sub> |
| 3DHSA | 3-dehydroshikimic acid | C<sub>7</sub>H<sub>8</sub>O<sub>5</sub> |
| 4HB | 4-hydroxybutyric acid | C<sub>4</sub>H<sub>8</sub>O<sub>3</sub> |
| 6PG | 6-phosphogluconic acid | C<sub>6</sub>H<sub>13</sub>O<sub>10</sub>P |
| ACO | aconitic acid | C<sub>6</sub>H<sub>6</sub>O<sub>6</sub> |
| ACoA | acetyl coenzyme A | C<sub>23</sub>H<sub>38</sub>N<sub>7</sub>O<sub>17</sub>P<sub>3</sub>S |
| ADI | adipic acid | C<sub>6</sub>H<sub>10</sub>O<sub>4</sub> |
| ADP | adenosine diphosphate | C<sub>10</sub>H<sub>15</sub>N<sub>5</sub>O<sub>10</sub>P<sub>2</sub> |
| AMP | adenosine monophosphate | C<sub>10</sub>H<sub>14</sub>N<sub>5</sub>O<sub>7</sub>P |
| ANTH | anthranilic acid | C<sub>7</sub>H<sub>7</sub>NO<sub>2</sub> |
| ATP | adenosine triphosphate | C<sub>10</sub>H<sub>16</sub>N<sub>5</sub>O<sub>13</sub>P<sub>3</sub> |
| Cre PO4 | creatine phosphate | C<sub>4</sub>H<sub>10</sub>N<sub>3</sub>O<sub>5</sub>P |
| CMP | cytidine monophosphate | C<sub>9</sub>H<sub>14</sub>N<sub>3</sub>O<sub>8</sub>P |
| CTP | cytidine triphosphate | C<sub>9</sub>H<sub>16</sub>N<sub>3</sub>O<sub>14</sub>P<sub>3</sub> |
| DHAP | dihydroxyacetone phosphate | C<sub>3</sub>H<sub>5</sub>O<sub>6</sub>P |
| E4P | D-erythrose 4-phosphate | C<sub>4</sub>H<sub>9</sub>O<sub>7</sub>P |
| F16DP | fructose 1,6-bisphosphate | C<sub>6</sub>H<sub>10</sub>O<sub>12</sub>P<sub>2</sub> |
| F6P | fructose 6-phosphate | C<sub>6</sub>H<sub>11</sub>O<sub>9</sub>P |
| FUM | fumaric acid | C<sub>4</sub>H<sub>4</sub>O<sub>4</sub> |
| G1P | glucose 1-phosphate | C<sub>6</sub>H<sub>13</sub>O<sub>9</sub>P |
| G6P | glucose 6-phosphate | C<sub>6</sub>H<sub>11</sub>O<sub>9</sub>P |
| GA3P | glyceraldehyde 3-phosphate | C<sub>3</sub>H<sub>5</sub>O<sub>6</sub>P |
| GDP | guanosine diphosphate | C<sub>10</sub>H<sub>15</sub>N<sub>5</sub>O<sub>11</sub>P<sub>2</sub> |
| Glyco | glycolic acid | C<sub>2</sub>H<sub>4</sub>O<sub>3</sub> |
| Glyox | glyoxylic acid | C<sub>2</sub>H<sub>2</sub>O<sub>3</sub> |
| GMP | guanosine monophosphate | C<sub>10</sub>H<sub>14</sub>N<sub>5</sub>O<sub>8</sub>P |
| GTP | guanosine triphosphate | C<sub>10</sub>H<sub>16</sub>N<sub>5</sub>O<sub>14</sub>P<sub>3</sub> |
| ISO | isocitric acid | C<sub>6</sub>H<sub>8</sub>O<sub>7</sub> |
| CIT | citric acid | C<sub>6</sub>H<sub>8</sub>O<sub>7</sub> |
| KGA | α-ketoglutaric acid | C<sub>5</sub>H<sub>6</sub>O<sub>5</sub> |
| LAC | lactic acid | C<sub>3</sub>H<sub>6</sub>O<sub>3</sub> |
| MAL | malic acid | C<sub>4</sub>H<sub>6</sub>O<sub>5</sub> |
| NAD | nicotinamide adenine dinucleotide (oxidised) | C<sub>21</sub>H<sub>27</sub>N<sub>7</sub>O<sub>14</sub>P<sub>2</sub> |
| NADH | nicotinamide adenine dinucleotide (reduced) | C<sub>21</sub>H<sub>29</sub>N<sub>7</sub>O<sub>14</sub>P<sub>2</sub> |
| NADP | nicotinamide adenine dinucleotide phosphate (oxidised) | C<sub>21</sub>H<sub>28</sub>N<sub>7</sub>O<sub>17</sub>P<sub>3</sub> |
| NADPH | nicotinamide adenine dinucleotide phosphate (reduced) | C<sub>21</sub>H<sub>30</sub>N<sub>7</sub>O<sub>17</sub>P<sub>3</sub> |
| OAA | oxaloacetic acid | C<sub>4</sub>H<sub>4</sub>O<sub>5</sub> |
| PABA | para-aminobenzoic acid | C<sub>7</sub>H<sub>7</sub>NO<sub>2</sub> |
| PEP | phosphoenolpyruvic acid | C<sub>3</sub>H<sub>5</sub>O<sub>6</sub>P |
| PPY | phenylpyruvic acid | C<sub>9</sub>H<sub>8</sub>O<sub>3</sub> |
| PYR | pyruvic acid | C<sub>3</sub>H<sub>4</sub>O<sub>3</sub> |
| R5P | D-ribose 5-phosphate | C<sub>5</sub>H<sub>11</sub>O<sub>8</sub>P |
| RL5P | D-ribulose 5-phosphate | C<sub>5</sub>H<sub>11</sub>O<sub>8</sub>P |
| S3P | shikimic acid 3-phosphate | $C_7H_{11}O_8P$ |
| SHIK | shikimic acid | $C_7H_{10}O_5$ |
| SUC | succinic acid | $C_4H_6O_4$ |
| UDP | uridine diphosphate | $C_9H_{14}N_2O_{12}P_2$ |
| UDPG | uridine diphosphate glucose | $C_{15}H_{24}N_2O_{17}P_2$ |
| UDPGA | uridine diphosphate glucuronic acid | $C_{15}H_{22}N_2O_{18}P_2$ |
| UDPNAG | uridine diphosphate N-acetylglucosamine | $C_{17}H_{27}N_3O_{17}P_2$ |
| UMP | uridine monophosphate | $C_9H_{13}N_2O_9P$ |
| UTP | uridine triphosphate | $C_9H_{15}N_2O_{15}P_3$ |
| X5P | D-xylulose 5-phosphate | $C_5H_{11}O_8P$ |

**Supplementary Table 5:** Results of Dunnett’s T3 and Holm-Šídák post-hoc tests for multiple comparisons.

| Measure | Comparison | t-score | Mean 1 | Mean 2 | SED | p-value |
| --- | --- | --- | --- | --- | --- | --- |
| *Experiment 1* |  |  |  |  |  |  |
| Dopamine (DA) content | Vehicle vs. IL-6 | 2.855 | 7.156 | 11.39 | 1.483 | 0.0325 |
|  | IL-6 vs. IL-6 + Melatonin | 2.525 | 11.39 | 7.645 | 1.483 | 0.0431 |
| HVA/DA ratio | Vehicle vs. IL-6 | 2.637 | 0.1146 | 0.06299 | 0.0196 | 0.0510 |
|  | Vehicle vs. Melatonin | 2.515 | 0.1146 | 0.06746 | 0.0188 | 0.0510 |
| SV2C | Vehicle vs. IL-6 | 2.377 | 0.4746 | 0.4199 | 0.0230 | 0.0804 |
| Reactive oxygen species (ROS) | Vehicle vs. Melatonin | 2.950 | 142533 | 140328 | 747.4 | 0.0250 |
| Total antioxidant capacity (TAC) | Vehicle vs. IL-6 | 3.430 | 9.595 | 9.853 | 0.0753 | 0.0168 |
|  | Vehicle vs. Melatonin | 1.842 | 9.595 | 9.733 | 0.0753 | 0.0926 |
|  | IL-6 vs. IL-6 + Melatonin | 2.660 | 9.853 | 9.668 | 0.0697 | 0.0438 |
| p-AMPK/AMPK ratio | Vehicle vs. IL-6 | 2.831 | 0.1416 | 0.1637 | 0.0078 | 0.0201 |
|  | Vehicle vs. Melatonin | 2.874 | 0.1416 | 0.1192 | 0.0078 | 0.0201 |
|  | IL-6 vs. IL-6 + Melatonin | 7.067 | 0.1637 | 0.1144 | 0.0070 | <0.0001 |
| p-Erk1/2/Erk1/2 ratio | Vehicle vs. IL-6 | 3.396 | 0.2903 | 0.3401 | 0.0147 | 0.0091 |
| p-GSK3β/GSK3β ratio | IL-6 vs. IL-6 + Melatonin | 3.056 | 0.3538 | 0.3133 | 0.0133 | 0.0203 |
| Glucose uptake | IL-6 vs. IL-6 + Melatonin | 2.631 | 29718 | 33845 | 1569 | 0.0473 |
| Spare respiratory capacity | Vehicle vs. IL-6 | 2.933 | 69.34 | 28.81 | 13.82 | 0.0420 |
|  | IL-6 vs. IL-6 + Melatonin | 3.109 | 28.81 | 67.20 | 12.35 | 0.0313 |
| Dihydroxyacetone phosphate (DHAP) | IL-6 vs. IL-6 + Melatonin | 2.697 | 12481 | 17771 | 1961 | 0.0411 |
| Glyceraldehyde 3-phosphate (GA3P) | IL-6 vs. IL-6 + Melatonin | 2.519 | 738.4 | 1039 | 119.3 | 0.0599 |
| Adenosine monophosphate (AMP) | Vehicle vs. IL-6 | 3.175 | 2032 | 3977 | 612.6 | 0.0149 |
|  | Vehicle vs. Melatonin | 2.702 | 2032 | 3688 | 612.6 | 0.0280 |
|  | IL-6 vs. IL-6 + Melatonin | 1.921 | 3977 | 5099 | 584.1 | 0.0699 |
| Adenosine diphosphate (ADP) | Vehicle vs. IL-6 | 3.896 | 29878 | 42208 | 3165 | 0.0029 |
|  | Vehicle vs. Melatonin | 3.248 | 29878 | 40157 | 3165 | 0.0084 |
| Cytidine monophosphate (CMP) | Vehicle vs. IL-6 | 2.521 | 234.4 | 414.4 | 71.40 | 0.0611 |
| Guanosine monophosphate (GMP) | Vehicle vs. IL-6 | 3.065 | 389.6 | 777.7 | 126.6 | 0.0190 |
|  | Vehicle vs. Melatonin | 2.784 | 389.6 | 742.1 | 126.6 | 0.0235 |
|  | IL-6 vs. IL-6 + Melatonin | 1.971 | 777.7 | 1016 | 120.7 | 0.0635 |
| Guanosine diphosphate (GDP) | Vehicle vs. IL-6 | 4.074 | 3037 | 4383 | 330.5 | 0.0019 |
|  | Vehicle vs. Melatonin | 3.276 | 3037 | 4119 | 330.5 | 0.0079 |
| Uridine monophosphate (UMP) | Vehicle vs. IL-6 | 3.069 | 738.7 | 1413 | 219.7 | 0.0188 |
|  | Vehicle vs. Melatonin | 2.648 | 738.7 | 1321 | 219.7 | 0.0315 |
| Uridine diphosphate (UDP) | Vehicle vs. IL-6 | 3.728 | 3588 | 5248 | 445.0 | 0.0043 |
|  | Vehicle vs. Melatonin | 3.081 | 3588 | 4959 | 445.0 | 0.0123 |
| Reduced nicotinamide adenine dinucleotide (NADH) | Vehicle vs. Melatonin | 3.244 | 870.4 | 3237 | 729.6 | 0.0090 |
|  | IL-6 vs. IL-6 + Melatonin | 3.727 | 1921 | 4640 | 729.6 | 0.0046 |
| **Experiment 2** |  |  |  |  |  |  |
| Superoxide (SOX) | Vehicle vs. 500μM DA | 7.882 | 14378 | 18051 | 466.0 | <0.0001 |
|  | 500μM DA vs. DA + Ascorbic acid | 7.727 | 18051 | 14451 | 466.0 | <0.0001 |
|  | DA + Melatonin vs. DA + Ascorbic acid | 8.237 | 18289 | 14451 | 466.0 | <0.0001 |
| Total antioxidant capacity (TAC) | Vehicle vs. 500μM DA | 2.343 | 9.705 | 9.886 | 0.0770 | 0.0997 |
|  | 500μM DA vs. DA + Ascorbic acid | 3.927 | 9.886 | 10.21 | 0.0832 | 0.0061 |
|  | DA + Melatonin vs. DA + Ascorbic acid | 4.632 | 9.827 | 10.21 | 0.0832 | 0.0019 |
| p-AMPK/AMPK ratio | 500μM DA vs. DA + Ascorbic acid | 3.090 | 0.1218 | 0.1494 | 0.0089 | 0.0248 |
| p-Erk1/2/Erk1/2 ratio | Vehicle vs. 500μM DA | 6.620 | 0.6013 | 0.9322 | 0.0500 | 0.0025 |
|  | 500μM DA vs. DA + Melatonin | 3.933 | 0.9322 | 0.7383 | 0.0493 | 0.0320 |
|  | 500μM DA vs. DA + Ascorbic acid | 8.843 | 0.9322 | 0.4918 | 0.0498 | 0.0005 |
|  | DA + Melatonin vs. DA + Ascorbic acid | 12.82 | 0.7383 | 0.4918 | 0.0192 | <0.0001 |
| p-mTOR/mTOR ratio | 500μM DA vs. DA + Ascorbic acid | 3.803 | 0.4935 | 0.3775 | 0.0305 | 0.0035 |
|  | DA + Melatonin vs. DA + Ascorbic acid | 4.215 | 0.5061 | 0.3775 | 0.0305 | 0.0015 |
| Insulin receptor alpha | Vehicle vs. 500μM DA | 3.827 | 0.0161 | 0.0285 | 0.0325 | 0.0360 |
| Glucose transporter 1 (GLUT1) | Vehicle vs. 500μM DA | 16.81 | 0.1981 | 0.5774 | 0.0226 | <0.0001 |
|  | 500μM DA vs. DA + Melatonin | 8.366 | 0.5774 | 0.3887 | 0.0226 | <0.0001 |
|  | 500μM DA vs. DA + Ascorbic acid | 17.08 | 0.5774 | 0.1920 | 0.0226 | <0.0001 |
|  | DA + Melatonin vs. DA + Ascorbic acid | 8.717 | 0.3887 | 0.1920 | 0.0226 | <0.0001 |
| Glucose uptake | 500μM DA vs. DA + Ascorbic acid | 3.336 | 28324 | 21847 | 1942 | 0.0110 |
|  | DA + Melatonin vs. DA + Ascorbic acid | 4.124 | 30247 | 21847 | 2037 | 0.0019 |
| Spare respiratory capacity | Vehicle vs. 500μM DA | 3.271 | 56.27 | 105.1 | 14.93 | 0.0103 |
|  | DA + Melatonin vs. DA + Ascorbic acid | 2.509 | 125.9 | 88.25 | 15.00 | 0.0616 |
| Maximal respiration | Vehicle vs. 500µM DA | 3.206 | 134.0 | 191.2 | 17.85 | 0.0123 |
| Basal glycolytic ATP production (GlycoATP) | Vehicle vs. 500µM DA | 3.175 | 193.3 | 269.4 | 23.97 | 0.0108 |
|  | 500µM DA vs. DA + Ascorbic acid | 2.935 | 269.4 | 197.2 | 24.62 | 0.0156 |
|  | DA + Melatonin vs. DA + Ascorbic acid | 3.706 | 286.4 | 197.2 | 24.09 | 0.0029 |
| Dihydroxyacetone phosphate (DHAP) | 500µM DA vs. DA + Ascorbic acid | 3.782 | 25051 | 37251 | 3225 | 0.0043 |
|  | DA + Melatonin vs. DA + Ascorbic acid | 2.691 | 28572 | 37251 | 3225 | 0.0492 |
| Fructose 1,6-bisphosphate (F16DP) | 500µM DA vs. DA + Ascorbic acid | 3.260 | 20958 | 29835 | 2723 | 0.0159 |
| Glyceraldehyde 3-phosphate (GA3P) | 500µM DA vs. DA + Ascorbic acid | 3.630 | 441.0 | 602.1 | 44.39 | 0.0063 |
|  | DA + Melatonin vs. DA + Ascorbic acid | 2.608 | 486.4 | 602.1 | 44.39 | 0.0592 |
| Glycerol-3-phosphate (G3P) | Vehicle vs. 500µM DA | 4.982 | 8248 | 4346 | 783.3 | 0.0035 |
|  | 500µM DA vs. DA + Ascorbic acid | 3.483 | 4346 | 6527 | 626.0 | 0.0268 |
| D-ribose 5-phosphate (R5P) | 500µM DA vs. DA + Ascorbic acid | 2.977 | 681.0 | 824.0 | 48.03 | 0.0296 |
|  | DA + Melatonin vs. DA + Ascorbic acid | 3.003 | 679.8 | 824.0 | 48.03 | 0.0296 |
| Sedoheptulose 7-phosphate (SH7P) | Vehicle vs. 500µM DA | 2.653 | 7726 | 4173 | 1340 | 0.0665 |
| Coenzyme A (CoA) | Vehicle vs. 500µM DA | 3.156 | 40973 | 24033 | 5367 | 0.0205 |
| Creatine phosphate (CrePO4) | 500µM DA vs. DA + Ascorbic acid | 4.694 | 1062635 | 750337 | 66531 | 0.0004 |
|  | DA + Melatonin vs. DA + Ascorbic acid | 3.303 | 970063 | 750337 | 66531 | 0.0115 |
| Oxidised nicotinamide adenine dinucleotide (NAD) | Vehicle vs. 5µM DA | 3.121 | 41500 | 36195 | 1700 | 0.0223 |
| Uridine diphosphate N-acetylglucosamine (UDPNAG) | 500µM DA vs. DA + Ascorbic acid | 4.110 | 50827 | 38969 | 2885 | 0.0015 |
|  | DA + Melatonin vs. DA + Ascorbic acid | 4.410 | 51693 | 38969 | 2885 | 0.0009 |

Abbreviations: SED = standard error of the difference between means.

**Supplementary Table 6:** Results of Dunn’s post-hoc test for multiple comparisons.

| **Measure** | **Comparison** | **z-score** | **Rank 1** | **Rank 2** | **p-value** |
| --- | --- | --- | --- | --- | --- |
| Non-mitochondrial oxygen consumption | IL-6 vs. IL-6 + Melatonin | 3.236 | 25.00 | 8.889 | 0.0036 |
|  | Vehicle vs. 500μM DA | 3.022 | 12.80 | 32.50 | 0.0126 |
| Glucose 1-phosphate (G1P) | Vehicle vs. Melatonin | 2.409 | 18.67 | 8.833 | 0.0480 |
| Acetyl coenzyme A (ACoA) | Vehicle vs. IL-6 | 2.449 | 17.17 | 7.167 | 0.0429 |
| Reactive oxygen species (ROS) | Vehicle vs. 500μM DA | 5.752 | 44.20 | 6.700 | <0.0001 |
|  | 500μM DA vs. DA + Ascorbic acid | 3.497 | 6.700 | 29.50 | 0.0023 |

## Notes

**Funding sources:** Funding for this project was provided by the University of Queensland. Heather K. Macpherson is supported by a Research Training Program stipend and tuition fee offset scholarship awarded by the University of Queensland.

### Competing Interest Statement

The authors have declared no competing interest.

## References

Abreu, T., & Braganca, M. (2015). The bipolarity of light and dark: A review on Bipolar Disorder and circadian cycles. J Affect Disord, 185, 219–229. doi:10.1016/j.jad.2015.07.017

Alexiuk, N. A., & Vriend, J. (1991). Effects of daily afternoon melatonin administration on monoamine accumulation in median eminence and striatum of ovariectomized hamsters receiving pargyline. Neuroendocrinology, 54(1), 55–61. Retrieved from https://www.ncbi.nlm.nih.gov/pubmed/1922678

Alexiuk, N. A., & Vriend, J. (2007). Melatonin: effects on dopaminergic and serotonergic neurons of the caudate nucleus of the striatum of male Syrian hamsters. J Neural Transm (Vienna), 114(5), 549–554. doi:10.1007/s00702-006-0582-7

Alghannam, A. F., Ghaith, M. M., & Alhussain, M. H. (2021). Regulation of Energy Substrate Metabolism in Endurance Exercise. Int J Environ Res Public Health, 18(9). doi:10.3390/ijerph18094963

Ali, T., & Kim, M. O. (2015). Melatonin ameliorates amyloid beta-induced memory deficits, tau hyperphosphorylation and neurodegeneration via PI3/Akt/GSk3beta pathway in the mouse hippocampus. J Pineal Res, 59(1), 47–59. doi:10.1111/jpi.12238

Ashok, A. H., Marques, T. R., Jauhar, S., Nour, M. M., Goodwin, G. M., Young, A. H., & Howes, O. D. (2017). The dopamine hypothesis of bipolar affective disorder: the state of the art and implications for treatment. Mol Psychiatry, 22(5), 666–679. doi:10.1038/mp.2017.16

Azzi, A. (2022). Oxidative Stress: What Is It? Can It Be Measured? Where Is It Located? Can It Be Good or Bad? Can It Be Prevented? Can It Be Cured? Antioxidants (Basel), 11(8). doi:10.3390/antiox11081431

Bakshi, V. P., & Kelley, A. E. (1991). Dopaminergic regulation of feeding behavior: I. Differential effects of haloperidol microinfusion into three striatal subregions. Psychobiology, 19(3), 223–232. 10.3758/BF03332072

Baldo, B. A., Sadeghian, K., Basso, A. M., & Kelley, A. E. (2002). Effects of selective dopamine D1 or D2 receptor blockade within nucleus accumbens subregions on ingestive behavior and associated motor activity. Behav Brain Res, 137(1-2), 165–177. doi:10.1016/s0166-4328(02)00293-0

Barbosa, I. G., Nogueira, C. R., Rocha, N. P., Queiroz, A. L., Vago, J. P., Tavares, L. P., … de Sousa, L. P. (2013). Altered intracellular signaling cascades in peripheral blood mononuclear cells from BD patients. J Psychiatr Res, 47(12), 1949–1954. doi:10.1016/j.jpsychires.2013.08.019

Barbosa-Mendez, S., Perez-Sanchez, G., & Salazar-Juarez, A. (2023). Agomelatine decreases cocaine-induced locomotor sensitisation and dopamine release in rats. World J Biol Psychiatry, 24(5), 400–413. doi:10.1080/15622975.2022.2123954

Barroso, E., Jurado-Aguilar, J., Wahli, W., Palomer, X., & Vazquez-Carrera, M. (2024). Increased hepatic gluconeogenesis and type 2 diabetes mellitus. Trends Endocrinol Metab, 35(12), 1062–1077. doi:10.1016/j.tem.2024.05.006

Berk, M., Dodd, S., Kauer-Sant’anna, M., Malhi, G. S., Bourin, M., Kapczinski, F., & Norman, T. (2007). Dopamine dysregulation syndrome: implications for a dopamine hypothesis of bipolar disorder. Acta Psychiatr Scand Suppl(434), 41–49. doi:10.1111/j.1600-0447.2007.01058.x

Bernier, L. P., York, E. M., & MacVicar, B. A. (2020). Immunometabolism in the Brain: How Metabolism Shapes Microglial Function. Trends Neurosci, 43(11), 854–869. doi:10.1016/j.tins.2020.08.008

Bersani, G., & Garavini, A. (2000). Melatonin add-on in manic patients with treatment resistant insomnia. Prog Neuropsychopharmacol Biol Psychiatry, 24(2), 185–191. doi:10.1016/s0278-5846(99)00097-4

Bradley, A. J., Webb-Mitchell, R., Hazu, A., Slater, N., Middleton, B., Gallagher, P., … Anderson, K. N. (2017). Sleep and circadian rhythm disturbance in bipolar disorder. Psychol Med, 47(9), 1678–1689. doi:10.1017/S0033291717000186

Brunoni, A. R., Supasitthumrong, T., Teixeira, A. L., Vieira, E. L., Gattaz, W. F., Bensenor, I. M., … Maes, M. (2020). Differences in the immune-inflammatory profiles of unipolar and bipolar depression. J Affect Disord, 262, 8–15. doi:10.1016/j.jad.2019.10.037

Capes-Davis, A., Theodosopoulos, G., Atkin, I., Drexler, H. G., Kohara, A., MacLeod, R. A., … Freshney, R. I. (2010). Check your cultures! A list of cross-contaminated or misidentified cell lines. Int J Cancer, 127(1), 1–8. doi:10.1002/ijc.25242

Capuron, L., Pagnoni, G., Drake, D. F., Woolwine, B. J., Spivey, J. R., Crowe, R. J., … Miller, A. H. (2012). Dopaminergic mechanisms of reduced basal ganglia responses to hedonic reward during interferon alfa administration. Arch Gen Psychiatry, 69(10), 1044–1053. doi:10.1001/archgenpsychiatry.2011.2094

Cardinali, D. P., & Vigo, D. E. (2017). Melatonin, mitochondria, and the metabolic syndrome. Cell Mol Life Sci, 74(21), 3941–3954. doi:10.1007/s00018-017-2611-0

Chacko, B. K., Kramer, P. A., Ravi, S., Benavides, G. A., Mitchell, T., Dranka, B. P., … Darley-Usmar, V. M. (2014). The Bioenergetic Health Index: a new concept in mitochondrial translational research. Clin Sci (Lond), 127(6), 367–373. doi:10.1042/CS20140101

Chitimus, D. M., Popescu, M. R., Voiculescu, S. E., Panaitescu, A. M., Pavel, B., Zagrean, L., & Zagrean, A. M. (2020). Melatonin’s Impact on Antioxidative and Anti-Inflammatory Reprogramming in Homeostasis and Disease. Biomolecules, 10(9). doi:10.3390/biom10091211

Cholico, G. N., Orlowska, K., Fling, R. R., Sink, W. J., Zacharewski, N. A., Fader, K. A., … Zacharewski, T. (2023). Consequences of reprogramming acetyl-CoA metabolism by 2,3,7,8-tetrachlorodibenzo-p-dioxin in the mouse liver. Sci Rep, 13(1), 4138. doi:10.1038/s41598-023-31087-9

Cipolla-Neto, J., Amaral, F. G., Afeche, S. C., Tan, D. X., & Reiter, R. J. (2014). Melatonin, energy metabolism, and obesity: a review. J Pineal Res, 56(4), 371–381. doi:10.1111/jpi.12137

Costa, M., Roman Meller, M., & Kapczinski, F. (2023). Bipolar disorder triggered by Covid-19 infection. Trends Psychiatry Psychother, 45, e20210430. doi:10.47626/2237-6089-2021-0430

Cuperfain, A. B., Kennedy, J. L., & Goncalves, V. F. (2020). Overlapping mechanisms linking insulin resistance with cognition and neuroprogression in bipolar disorder. Neurosci Biobehav Rev, 111, 125–134. doi:10.1016/j.neubiorev.2020.01.022

D’Imperio, A., Lo, J., Bettini, L., Prada, P., & Bondolfi, G. (2022). Bipolar type I diagnosis after a manic episode secondary to SARS-CoV-2 infection: A case report. Medicine (Baltimore), 101(31), e29633. doi:10.1097/MD.0000000000029633

Dashty, M. (2013). A quick look at biochemistry: carbohydrate metabolism. Clin Biochem, 46(15), 1339–1352. doi:10.1016/j.clinbiochem.2013.04.027

Dmitrzak-Weglarz, M., Banach, E., Bilska, K., Narozna, B., Szczepankiewicz, A., Reszka, E., … Pawlak, J. (2021). Molecular Regulation of the Melatonin Biosynthesis Pathway in Unipolar and Bipolar Depression. Front Pharmacol, 12, 666541. doi:10.3389/fphar.2021.666541

Emmons, H. A., Wallace, C. W., & Fordahl, S. C. (2023). Interleukin-6 and tumor necrosis factor-alpha attenuate dopamine release in mice fed a high-fat diet, but not medium or low-fat diets. Nutr Neurosci, 26(9), 864–874. doi:10.1080/1028415X.2022.2103613

Eslami Amirabadi, M. R., Rajezi Esfahani, S., Davari-Ashtiani, R., Khademi, M., Emamalizadeh, B., Movafagh, A., … Razjoyan, K. (2015). Monoamine oxidase a gene polymorphisms and bipolar disorder in Iranian population. Iran Red Crescent Med J, 17(2), e23095. doi:10.5812/ircmj.23095

Espinosa, M. I., Gonzalez-Garcia, R. A., Valgepea, K., Plan, M. R., Scott, C., Pretorius, I. S., … Williams, T. C. (2020). Adaptive laboratory evolution of native methanol assimilation in Saccharomyces cerevisiae. Nat Commun, 11(1), 5564. doi:10.1038/s41467-020-19390-9

Etain, B., Dumaine, A., Bellivier, F., Pagan, C., Francelle, L., Goubran-Botros, H., … Jamain, S. (2012). Genetic and functional abnormalities of the melatonin biosynthesis pathway in patients with bipolar disorder. Hum Mol Genet, 21(18), 4030–4037. doi:10.1093/hmg/dds227

Exposito, I., Mora, F., & Oaknin, S. (1995). Dopamine-glutamic acid interaction in the anterior hypothalamus: modulatory effect of melatonin. Neuroreport, 6(4), 661–665. doi:10.1097/00001756-199503000-00019

Exposito, I., Mora, F., Zisapel, N., & Oaknin, S. (1995). The modulatory effect of melatonin on the dopamine-glutamate interaction in the anterior hypothalamus during ageing. Neuroreport, 6(17), 2399–2403. doi:10.1097/00001756-199511270-00029

Fan, M., Liu, B., Jiang, T., Jiang, X., Zhao, H., & Zhang, J. (2010). Meta-analysis of the association between the monoamine oxidase-A gene and mood disorders. Psychiatr Genet, 20(1), 1–7. doi:10.1097/YPG.0b013e3283351112

Felger, J. C., Li, Z., Haroon, E., Woolwine, B. J., Jung, M. Y., Hu, X., & Miller, A. H. (2016). Inflammation is associated with decreased functional connectivity within corticostriatal reward circuitry in depression. Mol Psychiatry, 21(10), 1358–1365. doi:10.1038/mp.2015.168

Felger, J. C., & Treadway, M. T. (2017). Inflammation Effects on Motivation and Motor Activity: Role of Dopamine. Neuropsychopharmacology, 42(1), 216–241. doi:10.1038/npp.2016.143

Filonenko, V., & Gout, I. (2023). Discovery and functional characterisation of protein CoAlation and the antioxidant function of coenzyme A. BBA Adv, 3, 100075. doi:10.1016/j.bbadva.2023.100075

Fu, B., Yilin Yao, Heng, D., Li, N., Ma, X., Wang, Q., … Zhang, C. (2022). The Effect of Melatonin on OCT4 Expression and Granulosa Cell Growth in Female Mice. Reprod Sci, 29(10), 2810–2819. doi:10.1007/s43032-021-00783-0

Gaddi, A. V., Galuppo, P., & Yang, J. (2017). Creatine Phosphate Administration in Cell Energy Impairment Conditions: A Summary of Past and Present Research. Heart Lung Circ, 26(10), 1026–1035. doi:10.1016/j.hlc.2016.12.020

Garaude, J. (2019). Reprogramming of mitochondrial metabolism by innate immunity. Curr Opin Immunol, 56, 17–23. doi:10.1016/j.coi.2018.09.010

Geddes, J. R., & Miklowitz, D. J. (2013). Treatment of bipolar disorder. Lancet, 381(9878), 1672–1682. doi:10.1016/S0140-6736(13)60857-0

Gomez-Santos, C., Ferrer, I., Santidrian, A. F., Barrachina, M., Gil, J., & Ambrosio, S. (2003). Dopamine induces autophagic cell death and alpha-synuclein increase in human neuroblastoma SH-SY5Y cells. J Neurosci Res, 73(3), 341–350. doi:10.1002/jnr.10663

Haeusler, R. A., McGraw, T. E., & Accili, D. (2018). Biochemical and cellular properties of insulin receptor signalling. Nat Rev Mol Cell Biol, 19(1), 31–44. doi:10.1038/nrm.2017.89

Huang, L., Frampton, G., Rao, A., Zhang, K. S., Chen, W., Lai, J. M., … DeMorrow, S. (2012). Monoamine oxidase A expression is suppressed in human cholangiocarcinoma via coordinated epigenetic and IL-6-driven events. Lab Invest, 92(10), 1451–1460. doi:10.1038/labinvest.2012.110

Hussain, T., Tan, B., Yin, Y., Blachier, F., Tossou, M. C., & Rahu, N. (2016). Oxidative Stress and Inflammation: What Polyphenols Can Do for Us? Oxid Med Cell Longev, 2016, 7432797. doi:10.1155/2016/7432797

Jenkins, D. E., Sreenivasan, D., Carman, F., Samal, B., Eiden, L. E., & Bunn, S. J. (2016). Interleukin-6-mediated signaling in adrenal medullary chromaffin cells. J Neurochem, 139(6), 1138–1150. doi:10.1111/jnc.13870

Jiki, Z., Lecour, S., & Nduhirabandi, F. (2018). Cardiovascular Benefits of Dietary Melatonin: A Myth or a Reality? Front Physiol, 9, 528. doi:10.3389/fphys.2018.00528

Kageyama, Y., Okura, S., Sukigara, A., Matsunaga, A., Maekubo, K., Oue, T., … Inoue, K. (2025). The Association Among Bipolar Disorder, Mitochondrial Dysfunction, and Reactive Oxygen Species. Biomolecules, 15(3). doi:10.3390/biom15030383

Kennedy, S. H., Kutcher, S. P., Ralevski, E., & Brown, G. M. (1996). Nocturnal melatonin and 24-hour 6-sulphatoxymelatonin levels in various phases of bipolar affective disorder. Psychiatry Res, 63(2-3), 219–222. doi:10.1016/0165-1781(96)02910-1

Kim, Y., Vadodaria, K. C., Lenkei, Z., Kato, T., Gage, F. H., Marchetto, M. C., & Santos, R. (2019). Mitochondria, Metabolism, and Redox Mechanisms in Psychiatric Disorders. Antioxid Redox Signal, 31(4), 275–317. doi:10.1089/ars.2018.7606

Kim, Y. K., Jung, H. G., Myint, A. M., Kim, H., & Park, S. H. (2007). Imbalance between pro-inflammatory and anti-inflammatory cytokines in bipolar disorder. J Affect Disord, 104(1-3), 91–95. doi:10.1016/j.jad.2007.02.018

Kirsch, M., & De Groot, H. (2001). NAD(P)H, a directly operating antioxidant? FASEB J, 15(9), 1569–1574. doi:10.1096/fj.00-0823hyp

Koga, N., Ogura, J., Yoshida, F., Hattori, K., Hori, H., Aizawa, E., … Kunugi, H. (2019). Altered polyunsaturated fatty acid levels in relation to proinflammatory cytokines, fatty acid desaturase genotype, and diet in bipolar disorder. Transl Psychiatry, 9(1), 208. doi:10.1038/s41398-019-0536-0

Kovalevich, J., & Langford, D. (2013). Considerations for the use of SH-SY5Y neuroblastoma cells in neurobiology. Methods Mol Biol, 1078, 9–21. doi:10.1007/978-1-62703-640-5_2

Lai, C. T., & Yu, P. H. (1997). Dopamine- and L-beta-3,4-dihydroxyphenylalanine hydrochloride (L-Dopa)-induced cytotoxicity towards catecholaminergic neuroblastoma SH-SY5Y cells. Effects of oxidative stress and antioxidative factors. Biochem Pharmacol, 53(3), 363–372. doi:10.1016/s0006-2952(96)00731-9

Lam, R. W., Berkowitz, A. L., Berga, S. L., Clark, C. M., Kripke, D. F., & Gillin, J. C. (1990). Melatonin suppression in bipolar and unipolar mood disorders. Psychiatry Res, 33(2), 129–134. doi:10.1016/0165-1781(90)90066-e

Lewy, A. J., Nurnberger, J. I., Jr., Wehr, T. A., Pack, D., Becker, L. E., Powell, R. L., & Newsome, D. A. (1985). Supersensitivity to light: possible trait marker for manic-depressive illness. Am J Psychiatry, 142(6), 725–727. doi:10.1176/ajp.142.6.725

Lewy, A. J., Wehr, T. A., Goodwin, F. K., Newsome, D. A., & Rosenthal, N. E. (1981). Manic-depressive patients may be supersensitive to light. Lancet, 1(8216), 383–384. doi:10.1016/s0140-6736(81)91697-4

Li, R., Hou, J., Xu, Q., Liu, Q. J., Shen, Y. J., Rodin, G., & Li, M. (2012). High level interleukin-6 in the medium of human pancreatic cancer cell culture suppresses production of neurotransmitters by PC12 cell line. Metab Brain Dis, 27(1), 91–100. doi:10.1007/s11011-011-9270-x

Li, W., Knowlton, D., Woodward, W. R., & Habecker, B. A. (2003). Regulation of noradrenergic function by inflammatory cytokines and depolarization. J Neurochem, 86(3), 774–783. doi:10.1046/j.1471-4159.2003.01890.x

Li, Y. S., Ren, H. C., & Cao, J. H. (2022). Roles of Interleukin-6-mediated immunometabolic reprogramming in COVID-19 and other viral infection-associated diseases. Int Immunopharmacol, 110, 109005. doi:10.1016/j.intimp.2022.109005

Lin, S. C., & Hardie, D. G. (2018). AMPK: Sensing Glucose as well as Cellular Energy Status. Cell Metab, 27(2), 299–313. doi:10.1016/j.cmet.2017.10.009

Liu, Z., Huang, L., Luo, X. J., Wu, L., & Li, M. (2016). MAOA Variants and Genetic Susceptibility to Major Psychiatric Disorders. Mol Neurobiol, 53(7), 4319–4327. doi:10.1007/s12035-015-9374-0

Livianos, L., Sierra, P., Arques, S., Garcia, A., & Rojo, L. (2012). Is melatonin an adjunctive stabilizer? Psychiatry Clin Neurosci, 66(1), 82–83. doi:10.1111/j.1440-1819.2011.02288.x

Madireddy, S., & Madireddy, S. (2022). Therapeutic Interventions to Mitigate Mitochondrial Dysfunction and Oxidative Stress-Induced Damage in Patients with Bipolar Disorder. Int J Mol Sci, 23(3). doi:10.3390/ijms23031844

Majidpoor, J., & Mortezaee, K. (2022). Interleukin-6 in SARS-CoV-2 induced disease: Interactions and therapeutic applications. Biomed Pharmacother, 145, 112419. doi:10.1016/j.biopha.2021.112419

Marchetti, P., Fovez, Q., Germain, N., Khamari, R., & Kluza, J. (2020). Mitochondrial spare respiratory capacity: Mechanisms, regulation, and significance in non-transformed and cancer cells. FASEB J, 34(10), 13106–13124. doi:10.1096/fj.202000767R

Martin-Vega, A., & Cobb, M. H. (2025). ERK1/2-MAPK signaling: Metabolic, organellar, and cytoskeletal interactions. Curr Opin Cell Biol, 95, 102526. doi:10.1016/j.ceb.2025.102526

Martorana, A., & Koch, G. (2014). “Is dopamine involved in Alzheimer’s disease?”. Front Aging Neurosci, 6, 252. doi:10.3389/fnagi.2014.00252

Massari, F., Ciccarese, C., Santoni, M., Iacovelli, R., Mazzucchelli, R., Piva, F., … Montironi, R. (2016). Metabolic phenotype of bladder cancer. Cancer Treat Rev, 45, 46–57. doi:10.1016/j.ctrv.2016.03.005

Meiser, J., Weindl, D., & Hiller, K. (2013). Complexity of dopamine metabolism. Cell Commun Signal, 11(1), 34. doi:10.1186/1478-811X-11-34

Mihaylova, M. M., & Shaw, R. J. (2011). The AMPK signalling pathway coordinates cell growth, autophagy and metabolism. Nat Cell Biol, 13(9), 1016–1023. doi:10.1038/ncb2329

Miola, A., De Filippis, E., Veldic, M., Ho, A. M., Winham, S. J., Mendoza, M., … Cuellar-Barboza, A. B. (2022). The genetics of bipolar disorder with obesity and type 2 diabetes. J Affect Disord, 313, 222–231. doi:10.1016/j.jad.2022.06.084

Moghaddam, H. S., Bahmani, S., Bayanati, S., Mahdavinasa, M., Rezaei, F., & Akhondzadeh, S. (2020). Efficacy of melatonin as an adjunct in the treatment of acute mania: a double-blind and placebo-controlled trial. Int Clin Psychopharmacol, 35(2), 81–88. doi:10.1097/YIC.0000000000000298

Morris, G., Walder, K., McGee, S. L., Dean, O. M., Tye, S. J., Maes, M., & Berk, M. (2017). A model of the mitochondrial basis of bipolar disorder. Neurosci Biobehav Rev, 74(Pt A), 1–20. doi:10.1016/j.neubiorev.2017.01.014

Mracek, T., Drahota, Z., & Houstek, J. (2013). The function and the role of the mitochondrial glycerol-3-phosphate dehydrogenase in mammalian tissues. Biochim Biophys Acta, 1827(3), 401–410. doi:10.1016/j.bbabio.2012.11.014

Nathan, P. J., Burrows, G. D., & Norman, T. R. (1999). Melatonin sensitivity to dim white light in affective disorders. Neuropsychopharmacology, 21(3), 408–413. doi:10.1016/S0893-133X(99)00018-4

Nierenberg, A. A. (2009). Low-dose buspirone, melatonin and low-dose bupropion added to mood stabilizers for severe treatment-resistant bipolar depression. Psychother Psychosom, 78(6), 391–393. doi:10.1159/000235985

Nierenberg, A. A., Agustini, B., Kohler-Forsberg, O., Cusin, C., Katz, D., Sylvia, L. G., … Berk, M. (2023). Diagnosis and Treatment of Bipolar Disorder: A Review. JAMA, 330(14), 1370–1380. doi:10.1001/jama.2023.18588

Nurnberger, J. I., Jr., Adkins, S., Lahiri, D. K., Mayeda, A., Hu, K., Lewy, A., … Davis-Singh, D. (2000). Melatonin suppression by light in euthymic bipolar and unipolar patients. Arch Gen Psychiatry, 57(6), 572–579. doi:10.1001/archpsyc.57.6.572

Pandey, G. N., Ren, X., Rizavi, H. S., & Zhang, H. (2015). Abnormal gene expression of proinflammatory cytokines and their receptors in the lymphocytes of patients with bipolar disorder. Bipolar Disord, 17(6), 636–644. doi:10.1111/bdi.12320

Paneque, A., Fortus, H., Zheng, J., Werlen, G., & Jacinto, E. (2023). The Hexosamine Biosynthesis Pathway: Regulation and Function. Genes (Basel), 14(4). doi:10.3390/genes14040933

Papa, S., Choy, P. M., & Bubici, C. (2019). The ERK and JNK pathways in the regulation of metabolic reprogramming. Oncogene, 38(13), 2223–2240. doi:10.1038/s41388-018-0582-8

Park, G., Jung, Y. S., Park, M. K., Yang, C. H., & Kim, Y. U. (2018). Melatonin inhibits attention-deficit/hyperactivity disorder caused by atopic dermatitis-induced psychological stress in an NC/Nga atopic-like mouse model. Sci Rep, 8(1), 14981. doi:10.1038/s41598-018-33317-x

Pe, L. S., Pe, K. C. S., Panmanee, J., Govitrapong, P., Yang, J. L., & Mukda, S. (2025). Plausible therapeutic effects of melatonin and analogs in the dopamine-associated pathophysiology of bipolar disorder. J Psychiatr Res, 182, 13–20. doi:10.1016/j.jpsychires.2024.12.046

Quested, D. J., Gibson, J. C., Sharpley, A. L., Cordey, J. H., Economou, A., De Crescenzo, F., … Geddes, J. R. (2021). Melatonin In Acute Mania Investigation (MIAMI-UK). A randomized controlled trial of add-on melatonin in bipolar disorder. Bipolar Disord, 23(2), 176–185. doi:10.1111/bdi.12944

Ribeiro, R. F. N., Santos, M. R., Aquino, M., de Almeida, L. P., Cavadas, C., & Silva, M. M. C. (2025). The Therapeutic Potential of Melatonin and Its Novel Synthetic Analogs in Circadian Rhythm Sleep Disorders, Inflammation-Associated Pathologies, and Neurodegenerative Diseases. Med Res Rev. doi:10.1002/med.22117

Robillard, R., Naismith, S. L., Rogers, N. L., Scott, E. M., Ip, T. K., Hermens, D. F., & Hickie, I. B. (2013). Sleep-wake cycle and melatonin rhythms in adolescents and young adults with mood disorders: comparison of unipolar and bipolar phenotypes. Eur Psychiatry, 28(7), 412–416. doi:10.1016/j.eurpsy.2013.04.001

Ruan, H., Li, X., Zhou, L., Zheng, Z., Hua, R., Wang, X., … Zheng, F. (2024). Melatonin decreases GSDME mediated mesothelial cell pyroptosis and prevents peritoneal fibrosis and ultrafiltration failure. Sci China Life Sci, 67(2), 360–378. doi:10.1007/s11427-022-2365-1

Saxton, R. A., & Sabatini, D. M. (2017). mTOR Signaling in Growth, Metabolism, and Disease. Cell, 168(6), 960–976. doi:10.1016/j.cell.2017.02.004

Schiller, E. D., Champney, T. H., Reiter, C. K., & Dohrman, D. P. (2003). Melatonin inhibition of nicotine-stimulated dopamine release in PC12 cells. Brain Res, 966(1), 95–102. doi:10.1016/s0006-8993(02)04200-2

Scott, M. R., & McClung, C. A. (2023). Bipolar Disorder. Curr Opin Neurobiol, 83. doi:10.1016/j.conb.2023.102801

Shieh, K. R., Chu, Y. S., & Pan, J. T. (1997). Circadian change of dopaminergic neuron activity: effects of constant light and melatonin. Neuroreport, 8(9-10), 2283–2287. doi:10.1097/00001756-199707070-00037

Silvestrini, A., & Mancini, A. (2024). The Double-Edged Sword of Total Antioxidant Capacity: Clinical Significance and Personal Experience. Antioxidants (Basel), 13(8). doi:10.3390/antiox13080933

Silvestrini, A., Meucci, E., Ricerca, B. M., & Mancini, A. (2023). Total Antioxidant Capacity: Biochemical Aspects and Clinical Significance. Int J Mol Sci, 24(13). doi:10.3390/ijms241310978

Sprenger, S., Bare, J. P., Kashyap, R., & Cardella, L. (2022). Acute mania following COVID-19 in a woman with no past psychiatric history case report. BMC Psychiatry, 22(1), 486. doi:10.1186/s12888-022-04110-y

Teixeira, A. L., Scholl, J. N., & Bauer, M. E. (2025). Psychoneuroimmunology of Mood Disorders. Methods Mol Biol, 2868, 49–72. doi:10.1007/978-1-0716-4200-9_4

Toyama, E. Q., Herzig, S., Courchet, J., Lewis, T. L., Jr., Loson, O. C., Hellberg, K., … Shaw, R. J. (2016). AMP-activated protein kinase mediates mitochondrial fission in response to energy stress. Science, 351(6270), 275–281. doi:10.1126/science.aab4138

Treadway, M. T., Cooper, J. A., & Miller, A. H. (2019). Can’t or Won’t? Immunometabolic Constraints on Dopaminergic Drive. Trends Cogn Sci, 23(5), 435–448. doi:10.1016/j.tics.2019.03.003

Uyanik, V., Tuglu, C., Gorgulu, Y., Kunduracilar, H., & Uyanik, M. S. (2015). Assessment of cytokine levels and hs-CRP in bipolar I disorder before and after treatment. Psychiatry Res, 228(3), 386–392. doi:10.1016/j.psychres.2015.05.078

van Niekerk, G., Christowitz, C., Conradie, D., & Engelbrecht, A. M. (2020). Insulin as an immunomodulatory hormone. Cytokine Growth Factor Rev, 52, 34–44. doi:10.1016/j.cytogfr.2019.11.006

Varela, R. B., Macpherson, H., Walker, A. J., Houghton, T., Yates, C., Yates, N. J., … Tye, S. J. (2025). Inflammation and metabolic dysfunction underly anhedonia-like behavior in antidepressant resistant male rats. Brain Behav Immun, 127, 170–182. doi:10.1016/j.bbi.2025.03.001

Vasconcelos-Moreno, M. P., Fries, G. R., Gubert, C., Dos Santos, B., Fijtman, A., Sartori, J., … Kauer-Sant’Anna, M. (2017). Telomere Length, Oxidative Stress, Inflammation and BDNF Levels in Siblings of Patients with Bipolar Disorder: Implications for Accelerated Cellular Aging. Int J Neuropsychopharmacol, 20(6), 445–454. doi:10.1093/ijnp/pyx001

Wang, L., Li, J., & Di, L. J. (2022). Glycogen synthesis and beyond, a comprehensive review of GSK3 as a key regulator of metabolic pathways and a therapeutic target for treating metabolic diseases. Med Res Rev, 42(2), 946–982. doi:10.1002/med.21867

Williams, A. C., & Ford, W. C. (2004). Functional significance of the pentose phosphate pathway and glutathione reductase in the antioxidant defenses of human sperm. Biol Reprod, 71(4), 1309–1316. doi:10.1095/biolreprod.104.028407

Winter, C., von Rumohr, A., Mundt, A., Petrus, D., Klein, J., Lee, T., … Juckel, G. (2007). Lesions of dopaminergic neurons in the substantia nigra pars compacta and in the ventral tegmental area enhance depressive-like behavior in rats. Behav Brain Res, 184(2), 133–141. doi:10.1016/j.bbr.2007.07.002

Won, E., & Kim, Y. K. (2017). An Oldie but Goodie: Lithium in the Treatment of Bipolar Disorder through Neuroprotective and Neurotrophic Mechanisms. Int J Mol Sci, 18(12). doi:10.3390/ijms18122679

Xicoy, H., Wieringa, B., & Martens, G. J. (2017). The SH-SY5Y cell line in Parkinson’s disease research: a systematic review. Mol Neurodegener, 12(1), 10. doi:10.1186/s13024-017-0149-0

Yang, S. Y., Hong, K. S., Cho, Y., Cho, E. Y., Choi, Y., Kim, Y., … Baek, J. H. (2021). Association between the Arylalkylamine N-Acetyltransferase (AANAT) Gene and Seasonality in Patients with Bipolar Disorder. Psychiatry Investig, 18(5), 453–462. doi:10.30773/pi.2020.0436

Young, L. T., Warsh, J. J., Kish, S. J., Shannak, K., & Hornykeiwicz, O. (1994). Reduced brain 5-HT and elevated NE turnover and metabolites in bipolar affective disorder. Biol Psychiatry, 35(2), 121–127. doi:10.1016/0006-3223(94)91201-7

Zalcman, S., Green-Johnson, J. M., Murray, L., Nance, D. M., Dyck, D., Anisman, H., & Greenberg, A. H. (1994). Cytokine-specific central monoamine alterations induced by interleukin-1, −2 and −6. Brain Res, 643(1-2), 40–49. doi:10.1016/0006-8993(94)90006-x

Zalcman, S., Murray, L., Dyck, D. G., Greenberg, A. H., & Nance, D. M. (1998). Interleukin-2 and −6 induce behavioral-activating effects in mice. Brain Res, 811(1-2), 111–121. doi:10.1016/s0006-8993(98)00904-4

Zhang, Z. Z., Lee, E. E., Sudderth, J., Yue, Y., Zia, A., Glass, D., … Wang, R. C. (2016). Glutathione Depletion, Pentose Phosphate Pathway Activation, and Hemolysis in Erythrocytes Protecting Cancer Cells from Vitamin C-induced Oxidative Stress. J Biol Chem, 291(44), 22861–22867. doi:10.1074/jbc.C116.748848

Zisapel, N., Egozi, Y., & Laudon, M. (1982). Inhibition of dopamine release by melatonin: regional distribution in the rat brain. Brain Res, 246(1), 161–163. doi:10.1016/0006-8993(82)90157-3

Zisapel, N., Egozi, Y., & Laudon, M. (1983). Inhibition by melatonin of dopamine release from rat hypothalamus in vitro: variations with sex and the estrous cycle. Neuroendocrinology, 37(1), 41–47. doi:10.1159/000123513

Zisapel, N., Egozi, Y., & Laudon, M. (1985). Circadian variations in the inhibition of dopamine release from adult and newborn rat hypothalamus by melatonin. Neuroendocrinology, 40(2), 102–108. doi:10.1159/000124062

Zisapel, N., & Laudon, M. (1982). Dopamine release induced by electrical field stimulation of rat hypothalamus in vitro: inhibition by melatonin. Biochem Biophys Res Commun, 104(4), 1610–1616. doi:10.1016/0006-291x(82)91437-1

Zisapel, N., & Laudon, M. (1983). Inhibition by melatonin of dopamine release from rat hypothalamus: regulation of calcium entry. Brain Res, 272(2), 378–381. doi:10.1016/0006-8993(83)90588-7

Zisapel, N., & Laudon, M. (1987). A novel melatonin antagonist affects melatonin-mediated processes in vitro and in vivo. Eur J Pharmacol, 136(2), 259–260. doi:10.1016/0014-2999(87)90722-9

